# Structure-based discovery of inhibitors of the SARS-CoV-2 Nsp14 *N7*-methyltransferase

**DOI:** 10.1101/2023.01.12.523677

**Authors:** Isha Singh, Fengling Li, Elissa Fink, Irene Chau, Alice Li, Annía Rodriguez-Hernández, Isabella Glenn, Francisco J. Zapatero-Belinchón, Mario Rodriguez, Kanchan Devkota, Zhijie Deng, Kris White, Xiaobo Wan, Nataliya A. Tolmachova, Yurii S. Moroz, H. Ümit Kaniskan, Melanie Ott, Adolfo Gastía-Sastre, Jian Jin, Danica Galonić Fujimori, John J. Irwin, Masoud Vedadi, Brian K. Shoichet

## Abstract

An under-explored target for SARS-CoV-2 is non-structural protein 14 (Nsp14), a crucial enzyme for viral replication that catalyzes the methylation of *N7*-guanosine of the viral RNA at 5′-end; this enables the virus to evade the host immune response by mimicking the eukaryotic post-transcriptional modification mechanism. We sought new inhibitors of the S-adenosyl methionine (SAM)-dependent methyltransferase (MTase) activity of Nsp14 with three large library docking strategies. First, up to 1.1 billion make-on-demand (“tangible”) lead-like molecules were docked against the enzyme’s SAM site, seeking reversible inhibitors. On de novo synthesis and testing, three inhibitors emerged with IC_50_ values ranging from 6 to 43 μM, each with novel chemotypes. Structure-guided optimization and *in vitro* characterization supported their non-covalent mechanism. In a second strategy, docking a library of 16 million tangible fragments revealed nine new inhibitors with IC_50_ values ranging from 12 to 341 μM and ligand efficiencies from 0.29 to 0.42. In a third strategy, a newly created library of 25 million tangible, virtual electrophiles were docked to covalently modify Cys387 in the SAM binding site. Seven inhibitors emerged with IC_50_ values ranging from 3.2 to 39 μM, the most potent being a reversible aldehyde. Initial optimization of a second series yielded a 7 μM acrylamide inhibitor. Three inhibitors characteristic of the new series were tested for selectivity against 30 human protein and RNA MTases, with one showing partial selectivity and one showing high selectivity. Overall, 32 inhibitors encompassing eleven chemotypes had IC_50_ values <50 μM and 5 inhibitors in four chemotypes had IC_50_ values <10 μM. These molecules are among the first non-SAM-like inhibitors of Nsp14, providing multiple starting points for optimizing towards antiviral activity.

## INTRODUCTION

The Covid-19 pandemic has inspired a search for targets whose inhibition would combat the virus. Fruits of such efforts have been the development of Paxlovid ^1^, an inhibitor of the major protease (MPro) of SARS-CoV-2, of Mulprinavir ^2, 3^, a disruptor of viral RNA polymerization, and the introduction of Remdesevir ^4, 5^, an RNA-dependent RNA polymerase (RdRp) inhibitor first developed to treat Ebola virus. These targets are well-precedented in antiviral research, with successful drugs treating analogous enzymes for HIV, HCV, RSV, HBV, HCMV, HSV, HPV, and human influenza virus ^6-9^, among others, and these SARS-CoV-2 enzymes have been the focus of enormous efforts among many groups ^1, 4, 5, 10-13^. Other SARS-CoV-2 enzymes have attracted less attention, likely because there is less precedence for their targeting as antivirals. Nevertheless, enzymes like the macrodomain ^14, 15^ and the papain-like protease of Nsp3 ^16^, and the MTases Nsp10-Nsp16 complex ^17^ and Nsp14 play key roles in the virulence of SARS-CoV-2 ^18-21^. While they have little precedence as antiviral drug targets, they seem attractive as novel enzymes for antiviral drug discovery.

Among these, the Nsp14 SAM-dependent MTase seemed attractive. This enzyme catalyzes the methylation of the *N7* position of the terminal guanine of viral RNA, forming a cap-0 structure similar to those in eukaryotic mRNA, which are required for translation ^18, 22-26^. Subsequently, Nsp10-Nsp16 methylates the 2′-O of the cap ribose to form cap-1 on the 3′ end. The capping of viral RNA by Nsp14 evades the host innate immune response to viral RNA, while ensuring efficient ribosome binding and engagement of the host-translational complex. Deletion of Nsp14 is thought to eliminate viral virulence, confirming its importance and potential status as a SARS-CoV-2 drug target ^18, 19^.

If SAM-dependent MTases have little precedence in antiviral chemotherapy, they have long been targeted in cancer chemotherapy ^27-29^. The binding determinants of these enzymes have been explored, especially in the SAM site, and several inhibitory analogs of the co-factor are available ^30^. This has supported the determination of the crystal structure of Nsp14 complexed with s-adenosyl homocysteine (SAH) ^23^, along with other structures ^23, 31^, and the development of an enzyme inhibition assay ^32, 33^. The latter revealed relatively potent SAM-like inhibitors. However, most of these were relatively large and charged, likely reducing permeability and bioavailability, and as SAM analogs likely to have activities against other MTases, especially the human Class I MTases ^33^. This makes them problematic as leads to new antivirals.

Building off the structural and enzymatic work, we sought to discover novel scaffolds, dissimilar to that of the SAM-like inhibitors previously investigated, that would complement the Nsp14 structure with better physical properties than the SAM analogs. We adopted a structure-based docking approach, where large libraries of “tangible”, make-on-demand molecules were fit into the SAM binding site of Nsp14. Those that fit well ^34-38^ were prioritized for synthesis and testing. Three libraries were docked: one composed of up to 1.1 billion lead-like molecules ^39^, one composed of 16 million fragment-like molecules, and a newly-constructed library of 25 million electrophiles, which might covalently modify the active-site Cys387 (**Figure 1**). Inhibitors emerged from all three campaigns, and subsequent structure-based optimization led to several classes of low μM inhibitors. Methodologically, it was interesting that the fragment screen revealed perhaps the most diverse set of compounds, and the set most useful for characterizing the binding site. This has also been seen for other SARS-CoV-2 targets such as Mpro ^40^ and Mac1 ^14, 15^ and is a point to which we will return.

**Figure 1.**
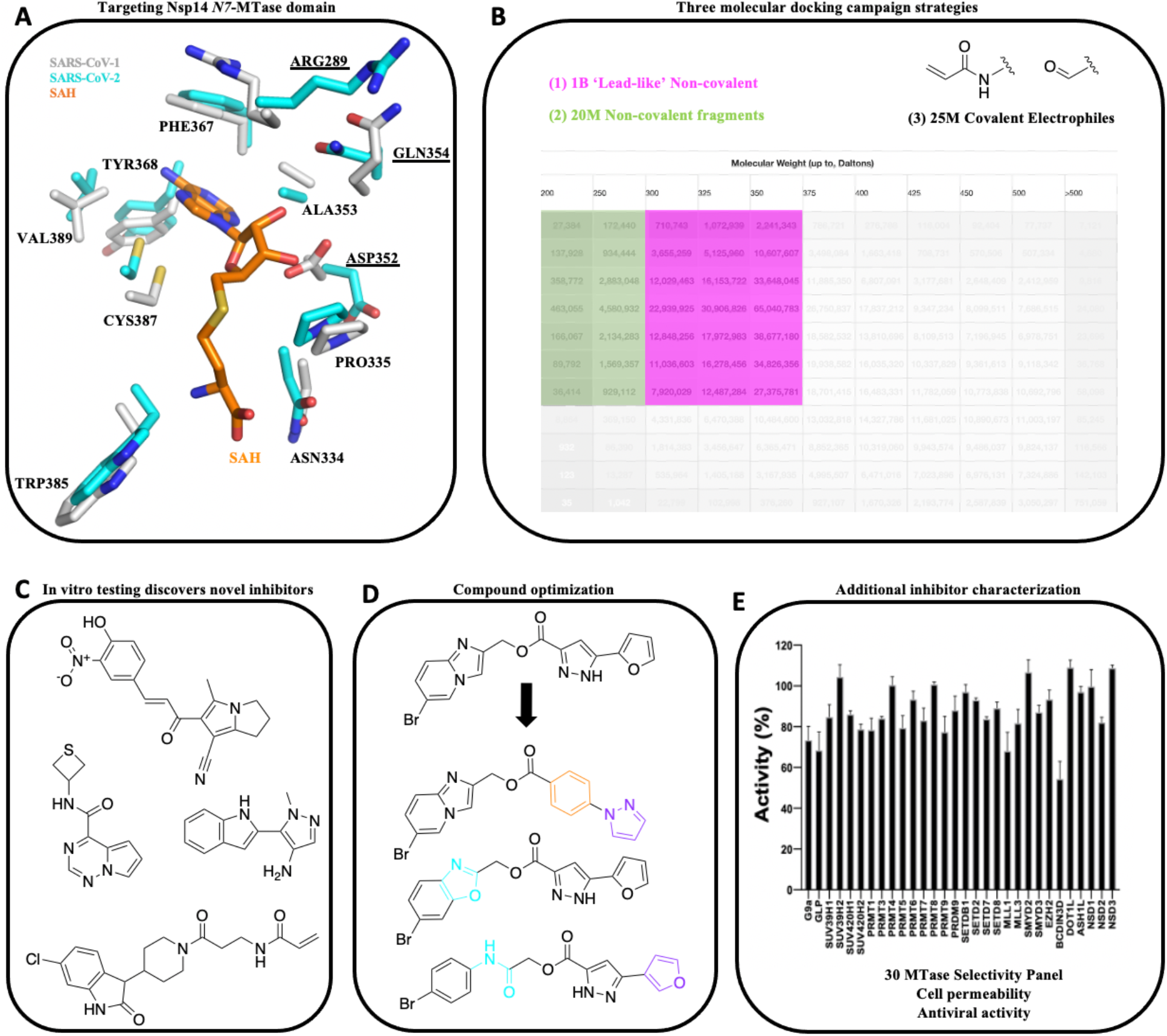
Workflow for inhibitor discovery against *N7-*MTase domain of Nsp14 using molecular docking. (**A**) SARS-CoV-1 and -2 Nsp14 MTase domains targeted with (**B**) three molecule subsets in molecular docking: lead-like non-covalent, fragment non-covalent, and acrylamide and aldehyde covalent electrophiles. (**C**) Diverse inhibitors discovered from each docking strategy followed by (**D**) compound optimization to improve potencies. (**E**) Best inhibitors evaluated for additional properties including MTase selectivity and antiviral efficacy.

## RESULTS

### Ultra-large library docking against Nsp14 identifies novel inhibitors

Due to the lack of SARS-CoV-2 Nsp14 protein structure when this project began, we initially used the *N7*-MTase domain of SARS-CoV-1 Nsp14 (PDB ID 5C8S) ^23^ for retrospective control calculations, which helped us to validate the recognition of known ligands. The SAM binding site of SARS-CoV-1 Nsp14 was used without any modifications as the active site residues are conserved in both SARS-CoV-1 and SARS-CoV-2 *N7*-MTase domains (**Figure 1A**). These control calculations confirmed that we could preferentially dock known MTase adenosyl-containing compounds (SAM, SAH, and Sinefungin) and other known MTase inhibitors including LLY283 ^41^, BMS-compd7f ^42^, and Epz04777 ^41-45^ in favorable geometries with high ranks versus 300 property matched decoys ^46, 47^. The structure used in these retrospective control calculations was subsequently supported by the recently-determined SAM binding pocket of *N7*-MTase domain of Nsp14 of SARS-CoV-2 (PDB ID 7N0B) ^31^; the C_*α*_ carbons of the two domains superpose with RMSD of 0.938 Å.

Seeking non-covalent inhibitors, we first docked over 680 million molecules, mostly in the “lead-like” range of the ZINC20 database (e.g., molecular weight ≤ 350 amu, cLogP values ≤ 3.5). Each library molecule was sampled for complementarity, in an average of 3438 orientations and for each of these about 187 conformations—over 3.6 × 10^14^ ligand configurations were sampled in the site in 121,018 core hours (about 5 days on 1000 cores). Seeking novel chemotypes, molecules topologically similar to SAM analogs were discarded. Compounds remaining were clustered based on ECFP4 fingerprints to identify unique chemotypes. Most cluster representatives were prioritized for interactions with Trp292, Gly333, Asn334, Asp352, Ala353, Phe367, Tyr368 and Val389 using LUNA ^48^. Molecules with strained conformations were deprioritized ^38^. Of the remaining molecules, the best scoring 5000 were visually inspected for key interactions and for unfavorable features, such as uncomplemented polar groups buried in the active site, using Chimera ^49^. Ultimately, 93 molecules, each in a different scaffold, were de novo synthesized and tested for enzyme inhibition at 30 and 50 μM, measuring the transfer of [^3^H]-methyl from the SAM methyl donor onto the cap structure of an RNA substrate (GpppAC_4_) (**Table S1**). Of the 93 molecules tested, only **ZINC475239213** (‘**9213**) inhibited by more than 50% and was considered active. This molecule had an IC_50_ of 20 μM in concentration-response (**Figure 2A**, middle panel). In its docked pose, the base-like moiety of ‘**9213** hydrogen-bonds with backbone amides of Ala353, Phe367 and Tyr368, while more distal parts of the molecule hydrogen-bond with Gln354 and Lys336 (**Figure 2A**, right panel). Van der Waals and stacking interactions are also apparent in the docked pose; overall these interactions resemble those observed among SAM and established SAM-like inhibitors but are made with different ligand groups.

**Figure 2.**
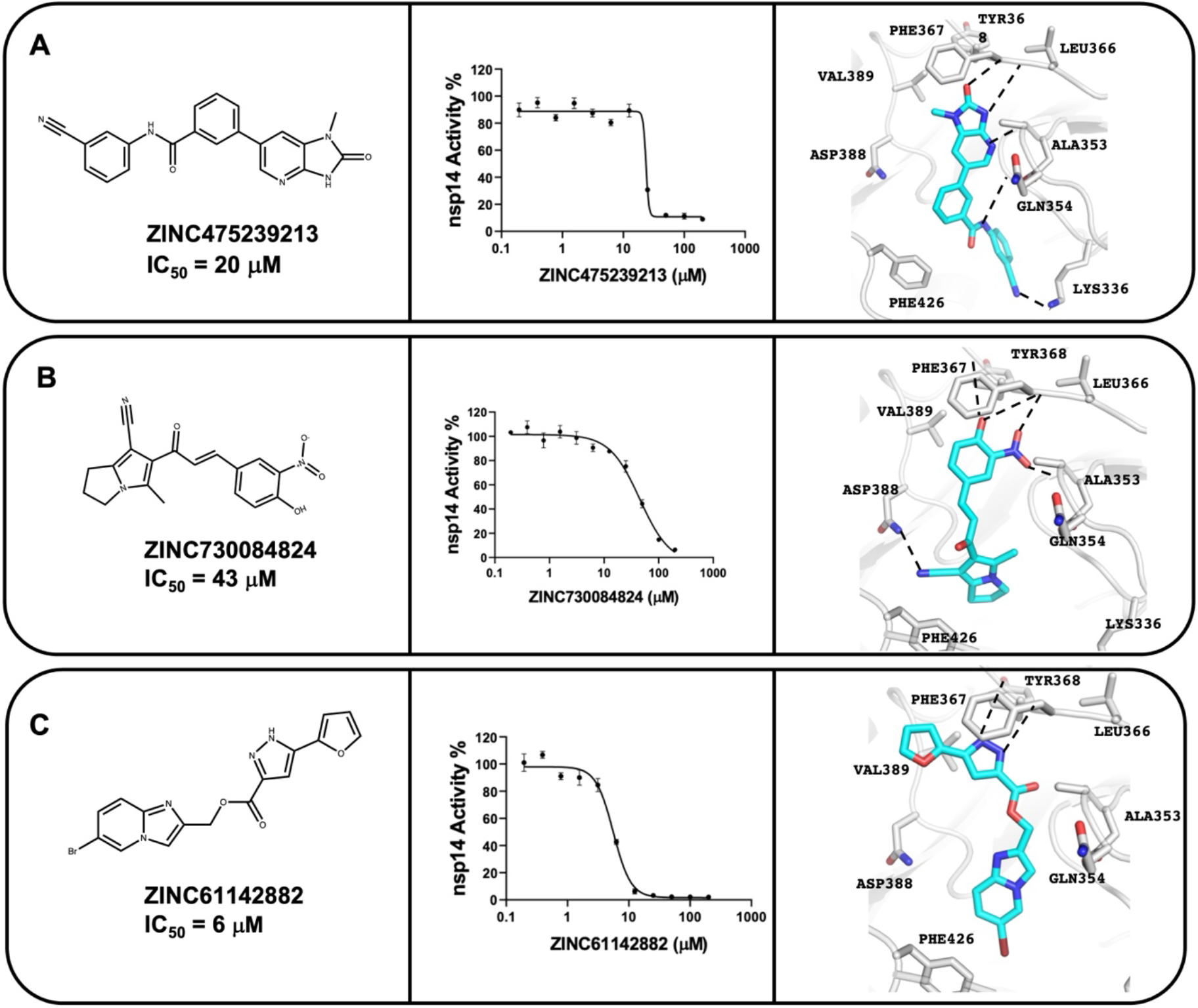
Ultra-large scale docking identifies three Nsp14 inhibitors with novel chemical scaffolds. 2D chemical structures, concentration-dependent Nsp14 MTase inhibition, and docked poses are represented for compounds ZINC475239213, ZINC730084824 and ZINC61142882 in panels **A, B** and **C**, respectively. SARS-CoV Nsp14 and inhibitors are shown in gray and cyan carbons, respectively, and hydrogen bonds are shown as black dashed lines. The experiments were performed in triplicate.

With the determination of the cryo-EM structure of the SARS-CoV-2 Nsp10-Nsp14 complex (PDB ID 7N0B) ^23^, and the development of a larger “tangible” ZINC22 ^50^ library of 1.1 billion molecules, we launched a second docking screen. The same retrospective control calculations were performed to optimize docking parameters, leading to similar sampling and calculation times. Following the same prioritization strategy as before, but seeking different chemotypes, 72 diverse molecules, were de novo synthesized and experimentally tested for enzyme inhibition (**Table S1**). Two inhibitors were found, **ZINC730084824** (‘**4824**) with an IC_50_ of 43 μM and **ZINC61142882** (‘**2882**) with an IC_50_ of 6 μM (**Figure 2B-C**), a hit rate of 2.7 %. The origins of the low hit rates for these two initial screens, and strategies to improve upon them, will be considered below.

### Optimization of the lead-like compounds

To improve the affinity of the three docking actives, we sought analogs among the 20 billion tangible molecules that have been enumerated in a version of the REAL database (http://enamine.net/compound-collections/real-compounds/real-space-navigator), using substructure and similarity searches in the SmallWorld (http://sw.docking.org) and Arthor (http://arthor.docking.org) search engines (NextMove Software, Cambridge UK) ^51^. Conservative analogs were prioritized, their structures and physical properties were calculated, and these were then docked into the Nsp14 SAM site. Analogs that docked to interact via π-π stacking with Phe367 or Trp292 and that appeared to hydrogen-bond with Tyr368, Ala353, Asp352 and Asp388 were prioritized. Overall, 12, 20 and 36 analogs of ‘**9213**, ‘**4824** and ‘**2882**, respectively, were synthesized and tested for enzyme inhibition (**Table S1, Figure 3**). The affinities of one of the analogs of ‘**2882** was similar to the initial hit, with an IC_50_ of 2.2 μM for compound ‘**1988** (**Figure 3C, Table S2**). The docked pose of ‘**1988** suggests a new hydrogen bond with Asp352 while conserving previous hydrogen bonds with Tyr368 and Ala353 (**Figure 3C**). Two **‘9213** analogs, **Z5347169163 (‘9163**, IC_50_ 15 μM) and **ZINC001342858621** (**‘8621**, IC_50_ 19 μM), also had similar affinities as the lead molecule (**Figure 3A, Table S3**). For ‘**4824**, two-fold improvement was observed for analogs **ZINC000916131631** (**‘1631**, IC_50_ 25 μM) and **Z5347186947** (**‘6947**, IC_50_ 19 μM) (**Figure 3B, Table S4**). Even though improvements were modest, the SAR was revealing. While little improvement was seen over the parent ‘**4824** or **‘9213**, for instance, many of the analogs tested remained relatively potent, with IC_50_ values often below 40 μM (**Table S2, Table S3, Table S4**). Moreover, replacement of the Michael acceptor vinyl group of **‘4824** in analogs **‘6947** and **Z5347186943** (**‘6943**, IC_50_ 40 μM), and removing the nitrile in analogs **‘1631** and **‘6947**, left molecules that remained active. In **‘9213**, this was also the case with the removal of the nitrile group in analogs **‘9163** and **‘8621**. These analogs support the idea that docking hits **‘9213** and **‘4824** are acting via a non-covalent mechanism, as modeled. Assessment of reversibility by jump dilution also suggest that inhibition by **‘4824** is not due to the nitrile electrophile (**Supplementary Figure 1**). ‘**1988** also appeared to be a non-covalent inhibitor in rapid dilution experiments (**Supplementary Figure 1**). In addition, **‘1988** and **‘4824** showed SAM- and RNA-competitive pattern of inhibition (**Supplementary Figure 2**). Taken together, these results support the non-covalent nature of binding of these families of molecules.

**Figure 3.**
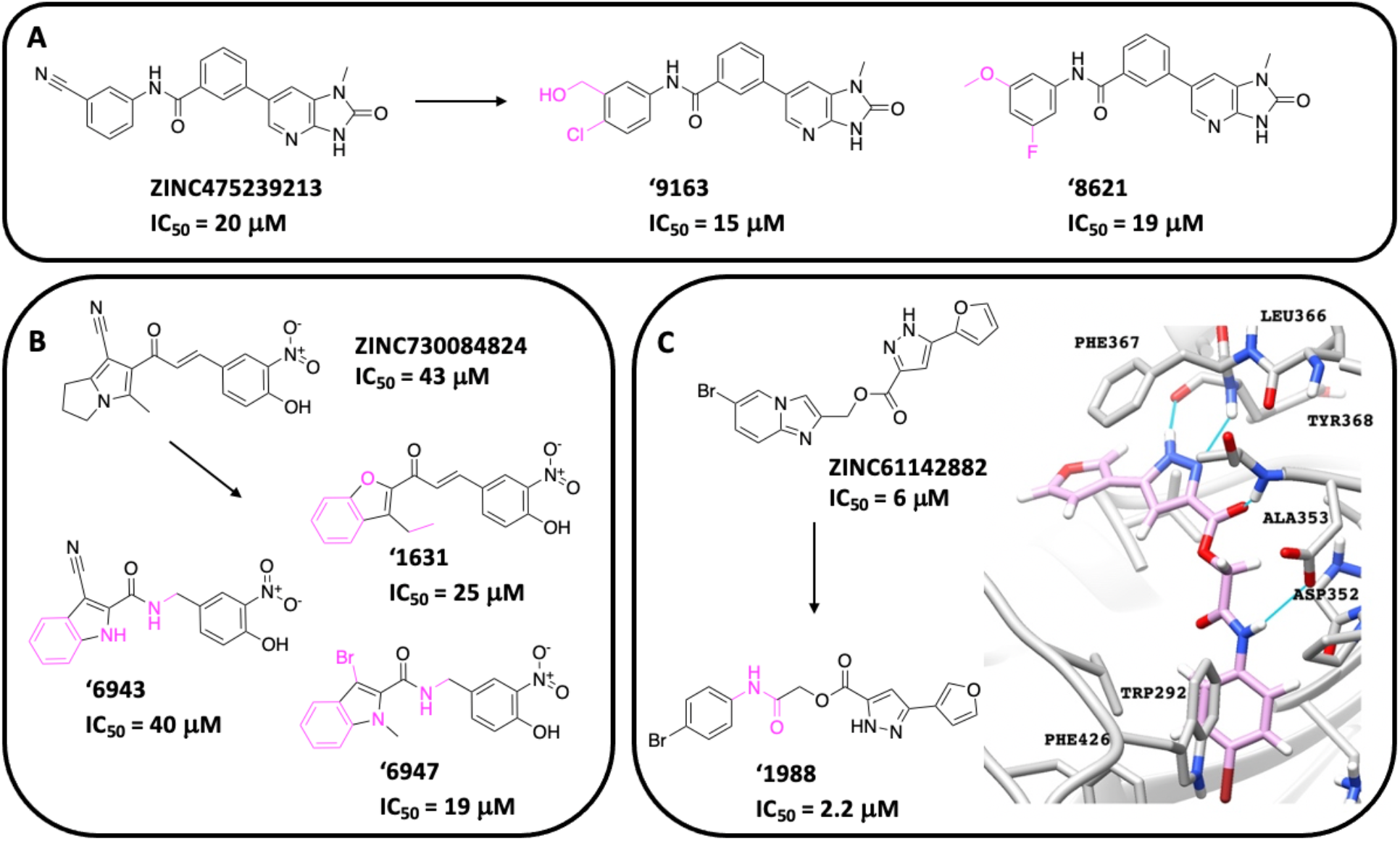
Hit optimization of the non-covalent compounds ‘9213, ‘4824, ‘2882. 2D chemical structures of the parent hit and corresponding analogs with chemical changes represented in pink. (**A**) ‘9213 analogs with the nitrile removed have similar IC_50_ values indicating non-covalent mechanism of action. (**B**) ‘4824 analogs with the nitrile or vinyl group removed have similar or more potent IC_50_ values. (**C**) The ‘**2882** analog **‘1988** is just as potent with opening of the bicyclic group. The **‘1988** docked pose (magenta carbons) is shown in SARS-CoV-2 Nsp14 (grey carbons). The experiments were performed in triplicate.

We tested the docking hits and analogs for colloidal aggregation, perhaps the dominant mechanism of artifactual activity in early discovery ^52-54^ (**Supplementary Figures 3.1** to **3.4**). As a first line of defense, all Nsp14 assays were conducted in the presence of 0.01% v/v Triton-X 100, a non-ionic detergent that disrupts colloidal aggregates and right-shifts their potency ^55, 56^. We also conducted follow-up assays for actives looking for particle formation by dynamic light scattering (DLS) and for activity against the widely-used counter-screening enzymes malate dehydrogenase (MDH) and AmpC β-lactamase (AmpC), both with and without detergent. The **‘9213** analog, ‘**3888**, did form colloid-like particles by DLS, with an apparent critical aggregation concentration (CAC) in the 10 uM range. Like most of the inhibitors studied here, however, the scattering intensity was relative modest, and the molecule did not inhibit the counter-screening enzymes at concentrations substantially higher than the IC_50_ for Nsp14, even the absence of detergent. While **‘3888** may form particles, we do not believe these are relevant for its inhibition of Nsp14. Compound ‘**4824** did not form detectable particles by DLS, but did inhibit the counter-screening enzymes MDH and AmpC at relevant concentrations. However, this inhibition can be at least partly attributed to strong absorbance at assay wavelengths (such absorbance was not an issue for the Nsp14 radioligand assay). Moreover, when we imitated the conditions of the Nsp14 assay by the addition of 0.01% Triton to the MDH and AmpC assays, inhibition was largely or entirely eliminated. Meanwhile, the ‘**4824** analogs (**‘6947**, ‘**6953**, ‘**6943**) did form colloid-like particles by DLS, though again with relatively modest scattering intensities. All three inhibited MDH or AmpC at relevant concentrations, but here too inhibition was largely eliminated by the addition of 0.01% of detergent. Compound ‘**1988** formed colloid-like particles by DLS, but with a CAC 10-fold higher than its Nsp14 IC_50_. While the compound’s inhibition of MDH was in a relevant range, it’s activity against AmpC was not, and the inhibition of both enzymes disappeared when we copied the Nsp14 conditions by the addition of Triton. We conclude that many of these inhibitors do aggregate, but this does not appear to be relevant for their inhibition of Nsp14, for which the inclusion of detergent appears to be prophylactic. These studies do support the usefulness of including detergents like Triton or Tween in enzyme and receptor inhibition assays.

### Docking 16 million fragment-like molecules

With only three inhibitor scaffolds discovered by lead-like docking, we stepped back to interrogate the site with fragment-based docking. Fragment screens explore more chemical space than a larger lead-like library ^14, 57, 58^, which may be helpful for an under-explored site where warheads and key residue interactions have not been characterized. With the proviso that they have lower affinities, fragments also have higher hit rates in empirical ^59^ and docking screens ^57, 58, 60^ than do lead-like molecules, providing a richer tiling of the binding site by ligand functional groups. Indeed, a strategy of fragment-docking was effective against another under-studied SARS-CoV-2 enzyme, Mac1 ^14, 15^, and fragment-based discovery nucleated a successful drug-discovery campaign against the Mpro enzyme ^10^. Accordingly, from the 16 million molecule fragment-like set (e.g., molecular weight ≤ 250, cLogP ≤ 2.5) in ZINC22, we targeted the full SARS-CoV-2 (PDB 7N0B) SAM site, the adenine portion of that site, and the SAM-tail region in three independent campaigns (**Figure 4A**) (Methods) ^61^. Overall, 14,406,946, 14,124,978, and 14,908,652 million molecules were scored, respectively. For each, the top-ranked 300,000 fragments were filtered as above, and the remaining fragments were clustered by topological similarity. Top-ranking cluster heads were visually inspected in Chimera ^49^ for favorable interactions, prioritizing those in the adenine site campaign for hydrogen bonds to Tyr368 and Ala353, and hydrophobic interactions with Phe367 ^48^. For the SAM-tail docking screen, interactions with Gly333 were prioritized, with additional interactions were selected for such as Gln313 and Asn386. For fragments docked against the entire SAM binding site, a combination of these interaction criteria were used. Ultimately 69 fragments were prioritized, of which 54 were successfully synthesized (78% fulfilment rate) (**Table S1**).

**Figure 4.**
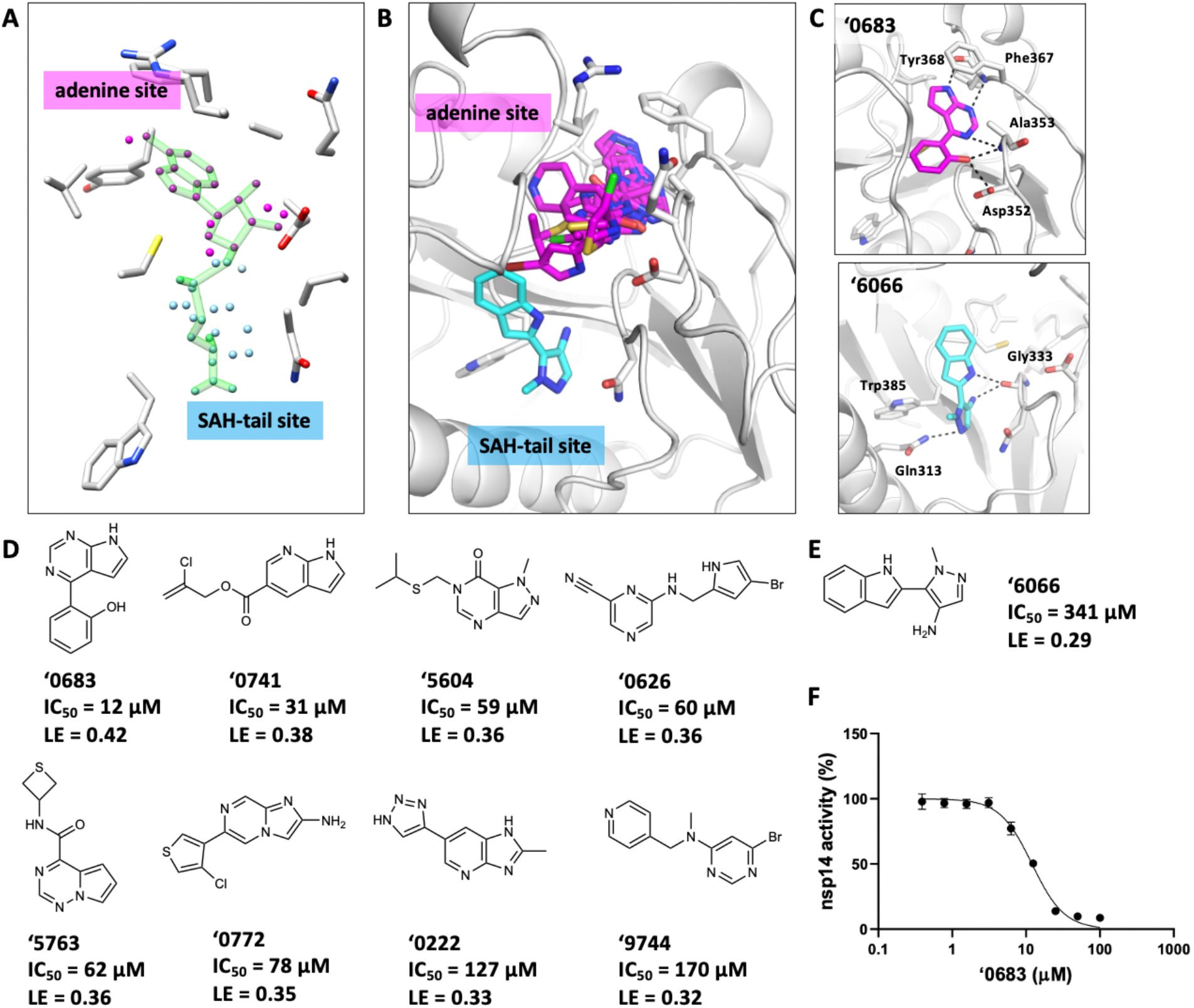
Fragment inhibitors from 16M docking screen. (**A**) Three sets of pseuodoatoms, which define where ligands are sampled in the binding site (“spheres”) used in the docking screens ^61^ including the adenine-site spheres (pink), SAM-tail site spheres (blue), and a superset of both. SARS-CoV-2 Nsp14 (grey carbons) with SAH (green carbons). (**B**) Overlay of all fragment docking hits in the SAM binding site of SARS-CoV-2 (tan carbons). Docked poses for adenine-site inhibitors shown (pink carbons) and SAM-tail site inhibitors (cyan carbons). (**C**) Docked poses of the best adenine-site fragment **‘0683**, and the SAM-tail site fragment **‘6066**. (**D**) Eight adenine-site fragment hits shown with their respective IC_50_ values. (**E**) The SAM-tail site fragment hit **‘6066**. (F) Concentration response curve of **‘0683** in the *N7*-MTase inhibitory activity assay. For D and E, IC_50_ values derived from concentration-response curves shown in Supplementary Figure 4. The experiments were performed in triplicate.

Of these 53, 9 fragments were hits with at least 50% inhibition at 300 μM and had IC_50_ values ranging from 12 μM to 341 μM (ligand efficiencies (LE) 0.29 to 0.42 kcal/heavy atoms). The most potent fragment, **‘0683**, had an IC_50_ of 12 μM and an LE of 0.42 kcal/heavy atom (**Figure 4, Supplementary Figure 4**). As with the larger lead-like inhibitors (above), **‘0683** was a competitive and presumably reversible inhibitor of both SAM and RNA binding (**Supplementary Figure 2**). In their docked poses, the fragment inhibitors were modeled to dock in the adenine-site forming hydrogen bonds with Tyr368, Phe367, or Ala353, often mimicking interactions of the adenine of SAM/SAH but with different functional groups and with diverse chemotypes (**Figure 4, Supplementary Figure 5**). One of the active fragments, **‘6066**, emerged from docking to the SAM-tail site, with hydrogen bonds to Gly333 and Gln313; we note that its activity was lower than most of those in the more tightly defined adenine site, with an IC_50_ of 341 μM (**Figure 4B, 4C, Figure 4E, Supplementary Figure 4**).

Fragment inhibitors were also evaluated for colloidal aggregation, as described above (**Supplementary Figure 3.1, 3.2, 3.3, 3.4**). While several did have CACs at relevant concentrations (**‘0222, ‘5604, 5763, 9744**), or inhibited MDH (**‘0741, ‘5604**) or both MDH and AmpC (**‘0683**) in the absence of detergent, most did not inhibit AmpC under any measured concentration, and on addition of detergent, mimicking the Nsp14 assay, MDH inhibition was largely or entirely eliminated (this effect was smaller for **‘0741**). Here too we believe that while under some conditions these fragments can form colloidal particles, such aggregation is not relevant for the Nsp14 inhibition we observe.

### Curation of 25 million aldehyde and acrylamide electrophiles for covalent docking

In a final strategy, we sought potential covalent electrophiles that could react with the enzyme’s active site Cys387. Such covalent docking has been successful in campaigns that targeted catalytic serine and non-catalytic, active site cysteine and lysine residues in enzymes such as β-lactamase, Jak kinases ^62^, eIF4e ^63^, M^Pro 64^ and targets such as RSK2 and MSK1 ^65^. These earlier campaigns had been limited to several hundred thousand electrophiles, largely from “in-stock” libraries. With the advent of the ultra-large tangible libraries, we thought to curate a larger set of electrophiles, focusing on aldehydes and acrylamides. Searching smarts patterns allowed us to build databases of 7.3 million aldehydes and 17.7 million acrylamides. We compared our aldehyde and acrylamide libraries to those that can be found in other in-stock or physical screening libraries, including the UCSF Small Molecule Discovery Center (SMDC) ^66^, Molecular Libraries Small Molecule Repository of the NIH (MLSMR) ^67^, and the in-stock set curated in ZINC20 ^51^. By total numbers, the aldehyde library is 196- to 10,000-fold larger than the number of aldehydes in the other libraries, while the number of core scaffolds represented by these electrophiles is 252- to 3,600-fold larger that those sampled in the previous libraries (**Table 1**). For acrylamides, there are 811- to 465,000-fold more electrophiles in the new library than in the public and in-stock sets, encompassing 250- to 58,000-fold more scaffolds. For the acrylamides, there are on average 13 molecules per scaffold in the new libraries, compared to 2-4 per scaffold in the other libraries. For aldehydes, the databases are comparable with our new library averaging 7 molecules per scaffold and 2-9 molecules per scaffold in the other databases. Given the rising interest in covalent-based inhibitors ^65, 68^, we have made these 25 million electrophiles openly available via http://covalent2022.docking.org in both 2D and DOCKovalent 3D format (Methods), along with ZINC and Enamine codes for ready acquisition.

**Table 1.** Expanded DOCKovalent electrophile databases.

| Database |  | Aldehydes |  | Acrylamides |  |
| --- | --- | --- | --- | --- | --- |
| Name | Size | # molecules | # BM Scaffolds | # molecules | # BM Scaffolds |
| Covalent2022.docking.org | 31,000,000,000 | 6,197,526 | 848,830 | 17,680,357 | 1,404,874 |
| ZINC20 In-Stock | 7,517,254 | 31,554 | 3,373 | 21,798 | 5,612 |
| UCSF SMDC | 690,125 | 615 | 235 | 38 | 24 |
| MLSMR | 406,098 | 908 | 407 | 63 | 27 |

### Covalent inhibitors from 25 million docking screen against Cys387

17.7 million acrylamides and 6.2 million aldehydes were docked against the SAM site adjacent to Cys387, using DOCKovalent ^62, 63^. Molecules were docked to form a covalent adduct with Cys387. Those with non-covalent DOCK3.7 scores < 0 kcal/mol were further filtered for internal strain ^38^, stranded hydrogen bond donors and acceptors, and for modeled hydrogen bonds with either Tyr368, Ala353, or Gly333 ^48^. Lastly, 33,156 molecules were clustered for topological similarity, and 9,591 molecules were prioritized for visual inspection in Chimera ^49^. From these, 92 molecules were selected for *de novo* synthesis. Of 61 aldehydes and 31 acrylamides, 47 and 26 were successfully synthesized, respectively, a 79% fulfilment rate (**Table S1**). On experimental testing, hits were defined as having at least 50% inhibition at 100 μM. For the aldehydes, four compounds were active of 51 tested (a hit rate of 8%) and had IC_50_ values ranging 3.2 to 19 μM (**Figure 5, Supplementary Figure 6, Supplementary Figure 7**). The most potent were **‘4975** and **‘1911** with IC_50_ values of 3.2 μM and 3.8, respectively (**Figure 5A**). These aldehydes were selected to hydrogen-bond with Ala 353 or Gly333 in their docked poses, and form additional hydrogen bonds in the pocket, including with Asn388, Arg310, Gln313, or Trp385 (**Figure 5D, Supplementary Figure 7**). Of the acrylamides, 2 of 26 tested had >50% inhibition at 100 μM, for a hit rate of 8%. (**Figure 5C, Supplementary Figure 6**). Inhibitor **acryl42** and **acryl41** had an IC_50_ values of 32 μM and 39 μM, respectively. **Acryl42** hydrogen bonds with Ala353 and Gly333 in its docked pose, while **acryl41** is a longer molecule extending farther than SAM/SAH, making hydrogen bonds with Gly333 and Gly313 in its docked pose (**Supplementary Figure 7**).

**Figure 5.**
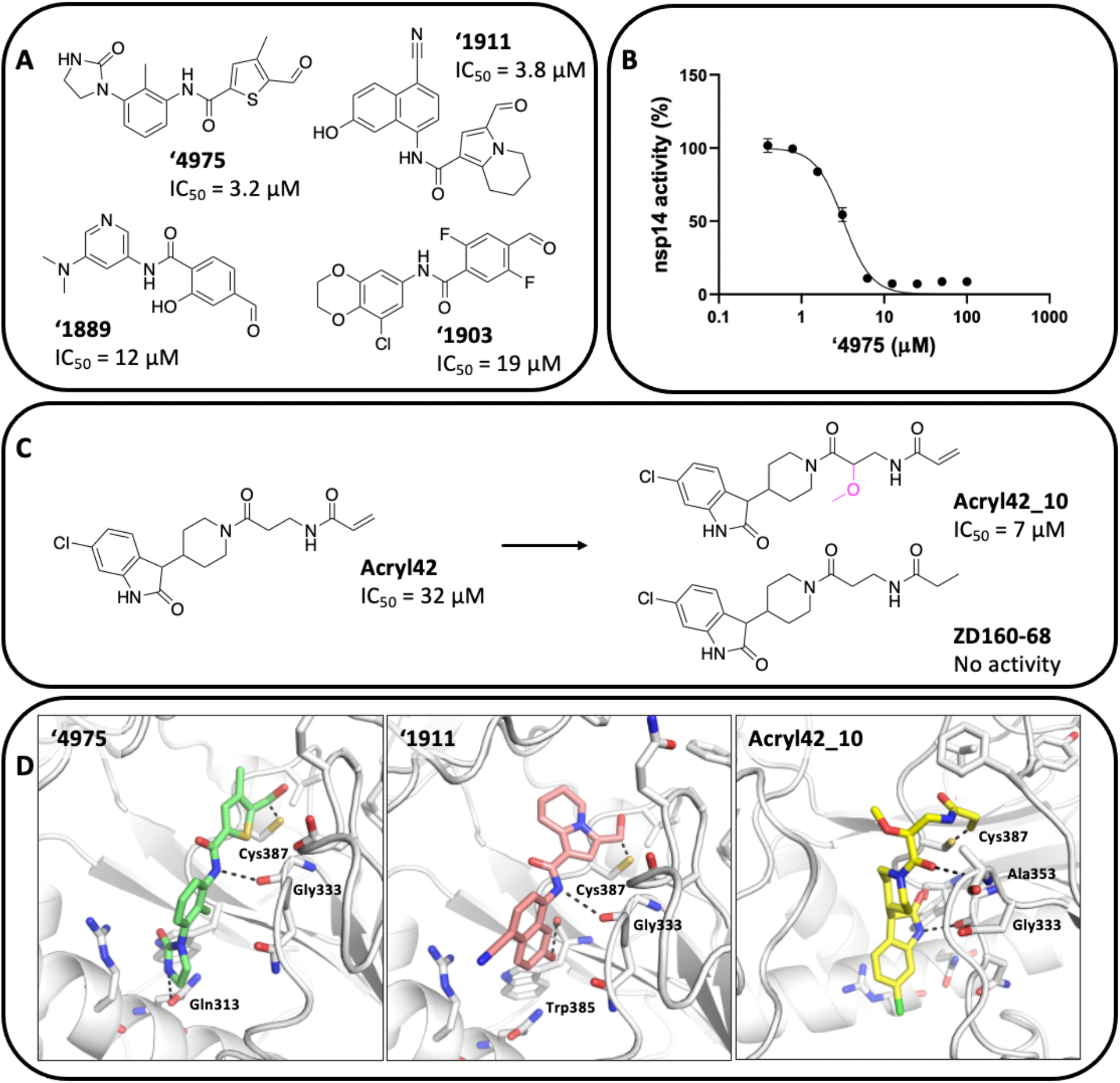
Docking 25 million electrophiles reveals aldehyde and acrylamide inhibitors. (**A**) Aldehyde docking hits. (**B**) Concentration-response curve for the most potent aldehyde **‘4975** in the *N7*-MTase inhibitory activity assay. (**C**) Acrylamide docking hits **acryl41** and **acryl42**, with analog **acryl42_10** and inactive analog **ZD160-68**. (**D**) Docked poses of **‘4975, ‘1911**, and modeled pose of analog **acryl42_10**. The experiments were performed in triplicate and additional concentration-response curves found in Supplementary Figure 6.

In early optimization of **acryl42**, analog **acryl42_10** was 4.5-fold more potent at 7 μM with the addition of a methoxy (**Figure 5C**). Adding a hydroxyl in the same place in analog **acryl42_11** resulted in an inactive analog, indicating the methoxy could be adding hydrophobic contacts, opposed to additional hydrogen bonds with the protein (**Table S6**). We tested the importance of the free amide of the acrylamide warhead with methylation of analog **acryl42_5**; the analog was inactive, perhaps reflecting the loss of a modeled hydrogen bond with the mainchain of Ala353.

We evaluated the covalent mechanism for the most potent covalent docking hits, aldehyde ‘**1911** and acrylamides **acryl41, acryl42**, and the **acryl42** analog, **acryl42_10**, first by mass spectrometry analysis. **‘1911** did not increase the molecular mass of Nsp14 (**Supplementary Figure 8**), likely reflecting the reversible binding of aldehydes to cysteines (Cys387 of Nsp14). In rapid dilution enzymatic experiments, **‘1911**, incubated at high concentrations, showed little residual inhibition when diluted below its IC_50_, further supporting a reversible covalent mechanism (**Supplementary Figure 1**). While **acryl41** did not form a measurable adduct by mass spectrometry, **acryl42** and its analog **acryl42_10** did do so, supporting a covalent inhibition mechanism (**Supplementary Figure 8**). We also changed the acrylamide warhead to the saturated propanamide group in compound **ZD160-68** resulting in no enzymatic inhibition, which furthered support for **acryl42** acting through covalent inhibition (**Figure 5C**). Overall, **acryl_42** and its analog, **acryl42_10**, appear to be irreversible covalent inhibitors, while **‘1911** appears to be a reversible covalent inhibitor. We expect that **acryl_41** is also acting as a covalent inhibitor but note that further mechanistic study of these classes is warranted.

The covalent inhibitors were evaluated for colloidal aggregation (**Supplementary Figure 3.1, 3.2, 3.3, 3.4**). The 12 uM aldehyde Z5185631889 (**‘1889** (**Figure 5**) had a CAC five-fold higher than its Nsp14 IC_50_ and did not inhibit either counter-screening enzyme under any measured condition—if this compound aggregates it is not relevant for its Nsp14 activity. While the 3.8 uM aldehyde **‘1911** (**Figure 5**) did form colloid-like particles by DLS at a relevant concentration, and did inhibit AmpC and MDH in the absence of Triton, this inhibition disappeared on the addition of the same amount of detergent (0.01% v/v) used in the Nsp14 assay. While this molecule likely is an aggregator, its aggregation is unlikely to be relevant to its Nsp14 inhibition.

### Selectivity against human protein and RNA methyltransferases

Three of the most potent inhibitors were counter-screened against a panel of 30 human SAM-dependent MTases. Compounds were tested for inhibition of the enzymes at 10 μM, then selected for IC_50_ determination if higher than 50% inhibition was observed. The non-covalent, 6 µM lead-like inhibitor, **‘1988**, showed only modest selectivity, inhibiting nine enzymes more than 50% with IC_50_ values ranging from 4 to 26 μM (**Figure 6**). The apparently reversible covalent, 8 µM **‘1911** had much better selectivity, inhibiting only two histone MTases G9a and G9a-like protein (GLP) with IC_50_ values of 30 μM and 14 μM, respectively. The most selective compound was **‘4824** with no inhibition greater than 50% for any enzyme in the panel.

**Figure 6.**
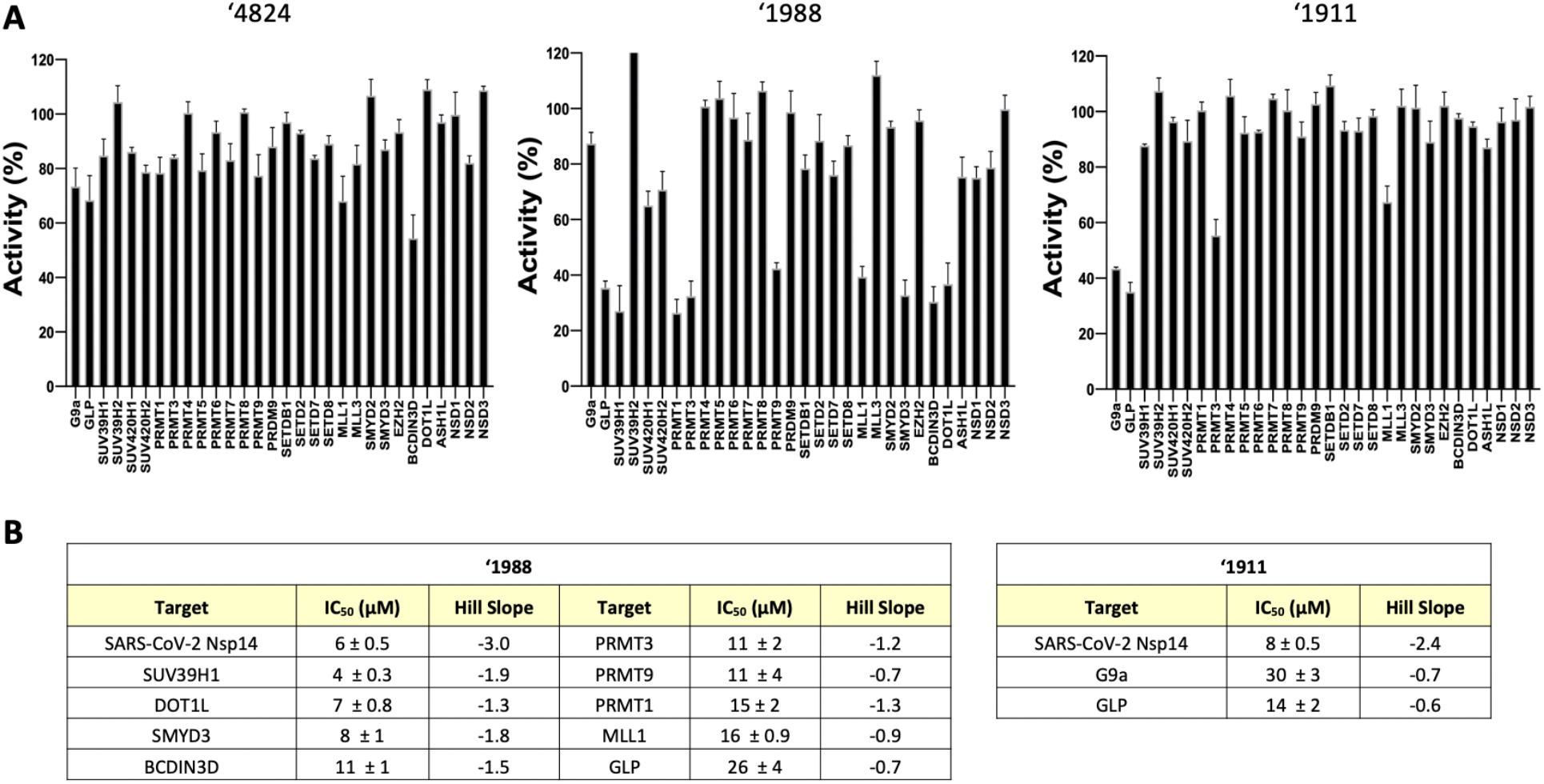
MTase selectivity of docking-derived inhibitors. (**A**) Compounds were tested against a panel of 30 SAM-dependent human protein and RNA MTases. Those with > 50% inhibition were prioritized for (**B**) IC_50_ determination. The experiments were performed in triplicate.

## DISCUSSION

From this study emerge among the first Nsp14 inhibitors unrelated to SAM, either topologically or by physical properties. Overall, 23 non-covalent, lead-like inhibitors across three scaffolds were found with IC_50_ values less than 50 μM, providing SAR for additional optimization (**Figure 2, Figure 3, Table S2, Table S3, Table S4**). Additional characterization and structure-based optimization demonstrated their competitive, non-covalent mechanism of action against Nsp14 (**Supplementary Figure 1, Supplementary Figure 2**). The most active covalent inhibitors were the initial aldehyde docking hits, with IC_50_ values ranging 3.5 to 12 μM, and the acrylamide analog **acryl42_10** with an IC_50_ of 7 μM, all modeled to modify Cys387 of Nsp14 (**Figure 5**). Finding these depended on developing new tangible libraries of 25 million electrophiles—these have been made publicly available for community use (https://covalent2022.docking.org) (**Table 1**). Another eight families of inhibitors were revealed from docking a library of 16 million tangible fragments (**Figure 4**). While affinities were naturally lower than the best of the lead-like inhibitors, several fragments had mid-μM IC_50_ values, and the four most potent had LEs 0.32 to 0.42 kcal/HAC. Taken together, 19 new chemotypes were found; of these, 11 had members with IC_50_ values <50 μM.

SARS-CoV-2 Nsp14 inhibitors described to date are SAM analogs ^33, 69, 70^ or fragments with extensive water networks ^71^. While the SAM analogs are widely-studied, they typically suffer from both low cell-permeability, owing to their size and ionization state, and from low selectivity, owing to their high similarity to the shared co-factor of this large family of MTases. Conversely, the new molecules described here are smaller and mostly uncharged, and topologically unrelated to SAM (**Table S7**). These properties may support optimization for cell permeability and bioavailability, and for selectivity. Consistent with this idea, in counter-screens of 30 SAM-dependent human protein and RNA MTases, **‘4824** was Nsp14 selective, and **‘1911** only hit two very closely related MTases (G9a and GLP).

Apart from the particular inhibitors discovered, lessons from this work may be useful for the other less-studied SARS-CoV-2 targets, which are the great majority of the viral genome. Unlike enzymes for which an investigational drug had been developed via similarities with other viruses, such as MPro or RdRp ^2-5, 72^, the MTase of Nsp14 had little inhibitor precedence on which to draw. Moreover, as a SAM-dependent enzyme with many related human enzymes, chemical novelty was important. Thus, as may be true with many SARS-CoV-2 targets, we could not leverage knowledge from previous chemical series other than SAM analogs. The lack of chemical precedence meant that these screens had a bootstrapping element to them—a small number of successes in early campaigns enabled us to optimize subsequent ones, contributing to improved hit-rates and affinities. We do note that our most informative screens—against the 16 million tangible fragments—occurred late in the campaign. Whereas there may still be skepticism about fragment docking, our own experience, not only here but also against the SARS-2 enzyme Mac1 _14, 15_ and in earlier studies against β-lactamases ^57, 58, 60^, is that fragment docking can reveal multiple chemotypes with high-ligand efficiency and fidelity to subsequently determined crystal structures. Were we to begin again, we might have started with the fragment screen, leveraging the interactions it revealed for campaigns against the larger, lead-like libraries. Such an approach may be useful against other understudied viral targets.

Certain caveats merit airing. Our most potent inhibitors are low-μM, weaker than the most potent of the SAM analogs previously characterized for Nsp14, the best of which inhibited in the 100 nM range ^33, 69, 70^. **‘1911** needs additional characterization of its reversible covalent mechanism of inhibition, limited here by its reversibility in mass spectrometry analysis, and low-μM activity in the rapid dilution experiments. Many of the inhibitors form colloidal aggregates, which would ordinarily be a concern for selectivity and artifactual activity. Control experiments suggest that such aggregation is not relevant for Nsp14 inhibition. Still, it remains true that this activity must be controlled for in subsequent optimization, and is a general hazard to navigation in early discovery. Importantly, antiviral activity, cell toxicity, and cell permeability remains to be explored for these molecules. Understanding these will inform future compound advancement.

These caveats should not obscure the key observation of this study, the discovery of nineteen new families of Nsp14 inhibitors. These new inhibitors are not only diverse, but they do not resemble the SAM-related molecules previously described for Nsp14 either topologically or by physical properties. They represent both non-covalent and covalent families, as well as fragments that tile the binding site. With the ongoing pandemic, they are being made openly available without restriction in the hopes that they may support a broad attack on this key but understudied target for antiviral drug discovery against Covid-19.

## Methods

### Non-covalent ultra-large scale docking

*N7*-MTase domain of SARS-CoV Nsp14 (PDB ID 5C8S) (12) and the *N7*-MTase domain of SARS-CoV2 Nsp14 from cryo-EM structure of the Nsp10-Nsp14 complex (PDB ID 7N0B) were used in two docking campaigns of >680 million “lead-like” molecules from the ZINC20 database (http://zinc20.docking.org) ^51^, and the ZINC22 >1.1 billion “lead-like” molecules (http://files.docking.org/zinc22), respectively, using DOCK3.7 ^37^. Forty-five matching spheres or local hot-spots generated from the crystal pose of SAM/SAH were used in the binding site for superimposing pre-generated flexible ligands and the different poses were scored by summing the different energies including; van der Waals interaction energies, Poisson-Boltzmann-based electrostatic interaction, and Generalized-Born/Surface Area-based ligand desolvation energies ^35, 36^. Receptor atoms were protonated with Reduce ^73^, and partial atomic charges were calculated using united-atom AMBER force field ^74^. AMBER atom-types were also used for Poisson-Boltzmann electrostatic potential energy grids using QNIFFT ^75^, CHEMGRID ^76^ was used for calculating van der Waals potential grids, and SOLVMAP ^35^ was used to calculate the Generalized-Born/Surface Area grids for ligand desolvation.

The docking setup was optimized for its ability to enrich knows MTase adenosyl group-containing compounds (SAM, SAH, and Sinefungin) and other known MTase inhibitors including Lly283, BMS-compd7f, and Epz04777 ^41-45^, in favorable geometries with high complementarity versus a set of property matched decoys ^46, 47^. About 50 decoys were generated for each ligand that had similar chemical properties to known ligands but were different topologically. The best optimized docking setup was evaluated for enrichment of ligands over decoys using log-adjusted area under the curve (logAUC values) ^46, 47^. All docked ligands were protonated with Marvin (version 15.11.23.0, ChemAxon, 2015; https://www.chemaxon.com) at pH 7.4, rendered into 3D with Corina (v.3.6.0026, Molecular Networks GmbH; https://www.mn-am.com/products/corina), and conformationally sampled using Omega (v.2.5.1.4, OpenEye Scientific Software; https://www.eyesopen.com/omega). Before docking the lead-like libraries, an ‘extrema set’ ^46, 77^ of 61,687 molecules was docked in the optimized system to ensure that the molecules with correct physical properties were enriched.

Overall, in the prospective screen, each library molecule was sampled in about 3438 orientations, on average about 187 conformations were sampled over five days on 1000 cores. The top-ranking 300,000 molecules were filtered for novelty using ECFP4-based Tanimoto coefficient (Tc <0.35) against known inhibitors of MTases. The remaining molecules were then clustered into related groups using an ECFP4-based Tc of 0.4. From the top 10,000 novel chemotypes, molecules with >2 kcal mol^−1^ internal strain ^38^ were excluded and the remaining candidates were visually inspected for best docked poses with favorable interactions with the SARS-CoV2 active site. Ultimately, overall 165 molecules were selected for de novo synthesis and testing.

### Non-covalent optimization

Analogs for docking hits **‘2882, ‘9213**, and **‘4824** were queried in Arthor and SmallWorld 1.4 and 12 billion make-on-demand libraries (http://sw.docking.org, http://arthor.docking.org), the latter primarily containing Enamine REAL compounds (http://enamine.net/compound-collections/real-compounds/real-space-navigator). The resulting analogs were further filtered based on Tc > 0.4 and docked to the *N7*-MTase domain of SARS-CoV2 Nsp14. Compounds were also designed by modifying 2D structure and custom synthesis by Enamine Ltd. (Kyïv, Ukraine). The docked poses were visually inspected for compatibility with the site and prioritized analogs were synthesized and tested for each series, respectively (**Table S1**).

### Fragment docking

The optimized docking setup from the SARS-CoV-2 second non-covalent lead-like screen described above was used. Three different screens were run with different matching spheres ^61^ – those in the adenine-site, SAM-tail site, or all matching spheres (**Figure 4A**), with 15,738,235 docked and 14,406,946 scored, 15,738,278 docked and 14,124,978 scored, and 16,299,173 docked and 14,908,652 scored, respectively. Each setup was analyzed separately until visualization in Chimera ^49^ – the top 300,000 ranked poses were filtered for having torsional strain less than 7 REU total, and single strain of 2.5 REU ^38^, less than 2 stranded hydrogen bond donors, less than 4 stranded hydrogen bond acceptors, and greater than 1 hydrogen bond to Tyr368, Ala353, or Gly333 ^48^. Remaining molecules were visually inspected for having favorable interactions. In total, 65 compounds were selected for purchasing, 50 from Enamine and 19 from WuXi, and overall, 53 were successfully synthesized for a fulfilment rate of 82%.

### Covalent database curation

SMARTS patterns for aldehydes or acrylamides ([CX3H1](=O)[#6] and ([CD1]=[CD2]-C(=O)-[NX3]), respectively) were searched in Enamine REAL databases, finding 20 million acrylamides and 6 million aldehydes. The DOCKovalent 3D files were generated as previously described ^63-65^. Briefly, the electrophiles were converted to their transition state product and a dummy atom was placed indicating to the docking algorithm which atom should be modeled covalently bound to the sulfur of the cysteine. Both 2D structures and 3D DOCKovalent files are now publicly available at http://covalent2022.docking.org. To compare to other public molecule databases, we used the ZINC20 in-stock set ^51^, the MLSMR library ^67^ and the UCSF SMDC library ^66^, and searched the same SMARTS patterns for acrylamides and aldehydes. The number of chemotypes were determined by Bemis-Murcko clustering ^78^.

### Covalent docking and compound optimization

The optimized docking setup from the first SARS-CoV-1 lead-like screen described above was used, with differences being which residues have been hyper-polarized ^77^ (Tyr368, Tyr368 and Ala353, or Tyr368, Ala353, and Gly333, referred to as 1-HP, 2-HP and 3-HP, respectively). For the acrylamide screen against 1-HP, molecules with docked scores less than 0 were selected for filtering (top 341,000); those with internal torsional strain less than total strain of 6.5 REU and single strain of 2 REU ^38^, molecules with less than 2 stranded hydrogen bond donors and less than 4 stranded hydrogen bond acceptors were prioritized. Molecules were also selected that formed at least one hydrogen bond to Tyr368, Ala353 or Gly333 using LUNA ^48^ leaving 2,423. After clustering for chemical similarity, 533 were visually inspected in Chimera ^49^. For the 2-HP setup, molecules with scores less than 0 (top 440,661) were filtered using the same criteria with 2,961 molecules remaining, comprising of 622 clusters that were visually inspected. For the 3-HP setup, no molecules passed the strain, IFP, and hydrogen bond filter and were not considered further. Visual inspection prioritized molecules with the same criterion as above. Lastly, selected compounds from both 1-HP and 2-HP setups were clustered to select unique chemotypes, and 31 were purchased. Synthesis was successful for 26 for a fulfilment rate of 84%.

For the aldehydes in the 1-HP setup, the top 894,979 compounds (dock score less than 0) were filtered to prioritized as the acrylamides were above, with clustering for chemical similarity leaving 1,340 for visual filtering. For the 2-HP setup, the top 1,494,350 were filtered to 3,548, and 3-HP setup of top 1,494,345 to 3,548 for visual inspection. Compounds were prioritized for the same interactions as the acrylamides, and finally 61 aldehydes were selected. Synthesis was successful for 47 of these for a fulfilment rate of 77%, and an overall covalent fulfillment rate of 79%.

Acryl42 analogs acryl42_5, acryl42_11 and acryl42_10 was designed off the acryl42 2D chemical structure, and synthesized by Enamine; ZD160-68 was designed to test the activity of the acrylamide warhead. The modeled pose of acryl42_10 was performed in Maestro (version 2021-2, Schrödinger, Inc.) by manually changing the acryl42 docked pose to acryl42_10 and minimizing the Nsp14-acryl42_10 complex using the Protein Preparation Wizard protocol.

### Make-on-demand synthesis

Compounds were purchased from Enamine Ltd. (Kyiv, Ukraine) or WuXi Appetec (Shanghai, China). Purities of active molecules were at least 90% and typically above 95% (based on LC/MS data) (**Table S1, Supplementary Figure 9**).

### Synthesis of ZD160-68

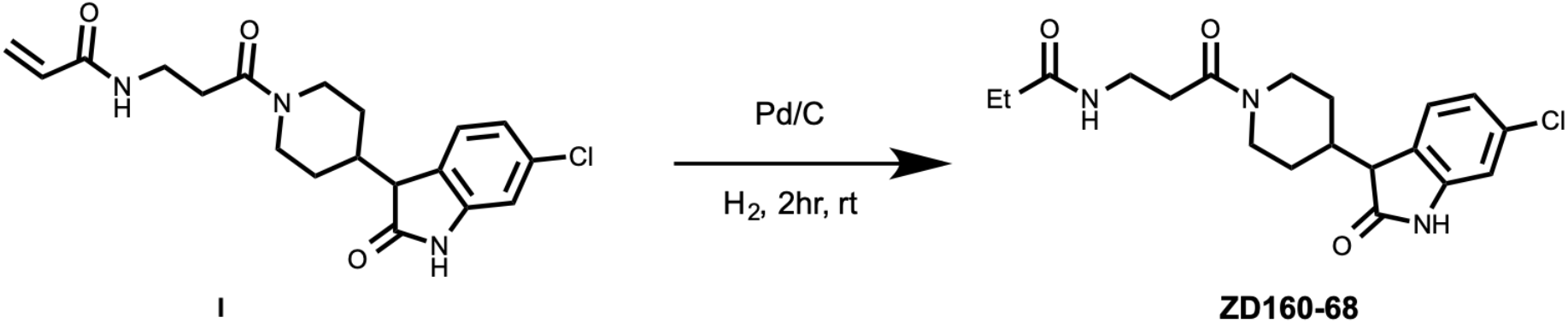

### N-(3-(4-(6-chloro-2-oxoindolin-3-yl)piperidin-1-yl)-3-oxopropyl)propionamide (ZD160-68)

To a solution of compound **I** (5.6 mg, 0.015 mmol) in THF (2 mL), was added Pd/C (10%, 5 mg). The mixture was stirred under H_2_ atmosphere for 2 hours followed by filtering. The filtrate was collected and purified by prep-HPLC to yield ZD160-68, white 1 solid (4 mg, 70% yield). ^**1**^**HNMR** (400 MHz, Methanol-*d*_4_) d 7.32 – 7.21 (m, 1H), 7.03 (dt, *J* = 8.0, 2.2 Hz, 1H), 6.91 (d, *J* = 2.2 Hz, 1H), 4.61 (t, *J* = 15.8 Hz, 1H), 4.00 (t, *J* = 13.9 Hz, 1H), 3.50 (q, *J* = 2.6 Hz, 1H), 3.42 (q, *J* = 6.3, 5.8 Hz, 2H), 3.11 – 3.00 (m, 1H), 2.63 – 2.52 (m, 3H), 2.36 (d, *J* = 12.7 Hz, 1H), 2.24 – 2.10 (m, 2H), 1.73 – 1.59 (m, 2H), 1.49 (dq, *J* = 17.2, 5.7, 4.3 Hz, 2H), 1.14 – 1.08 (m, 3H). **MS (ESI) *m/z*:** [M+H]^+^ calcd for C_19_H_24_ClN_3_O_3_ 378.8; found 378.3.

### Nsp14 expression and purification

For activity assays, Nsp14 was expressed and purified as previously described ^33^. The codon-optimized gene was also sub-cloned in a modified pET28b with 6x-Histidine and SUMO tag at the N-terminus. Nsp14 was expressed in *E. coli* Rosetta2(DE3) PlysS cells, growing in terrific broth at 37°C, induced at 18°C with 0.4 mM IPTG at OD 600nm of 1.2 for 18 h. Cell pellets were recovered and stored at -80°C.

For purification, cells were suspended in lysis buffer containing 50 mM HEPES, 500 mM NaCl, 10 mM imidazole, 10% v/v glycerol, 5 mM MgCl_2_, 1 mM TCEP pH 8.1 supplemented with EDTA-free protease cocktail inhibitor tablets (Thermo Scientific). Cells were disrupted by sonication and lysate centrifuged at 16,000 rpm for 30 min. Nsp14 was purified using a 5 mL HisTrap HP column, washed with 20 column volumes of lysis buffer with additional 20 and 30 mM imidazole, and eluted with buffer containing 500 mM imidazole. Protein fractions were exchanged to 50mM HEPES, 150 mM NaCl, 5 mM MgCl_2_, 10% glycerol, and 1 mM TCEP pH 8.0, and incubated overnight at 4°C with SenP1 protease at a 1:100 mass ratio. SUMO tag was removed using a MonoQ 10/100 column, pre-equilibrated in 50 mM HEPES, 20 mM NaCl, 5 mM MgCl_2_, and 1 mM TCEP pH 8.0. Nsp14 was in the unbound fraction. As a final step, the protein was purified using a size exclusion column s200 16/600 in the same buffer for SenP1 digestion. Purest fractions were pulled together, flash frozen, and stored at -80°C until needed.

### Enzyme Inhibition

The inhibitory effect of compounds on the methyltransferase activity of SARS-CoV-2 Nsp14 was assessed using a previously developed radiometric assay ^33^.

### Jump dilution

The recovery of Nsp14 activity after incubation with each inhibitor and rapid dilution was monitored. Nsp14 at 100-fold higher concentration than what required for activity measurement (1.5 nM) was incubated with each compound at 10-fold of IC_50_ value concentration for 1h at room temperature. Reaction mixtures were then rapidly diluted 100-fold into the assay buffer containing substrate RNA and SAM, and recovery of the Nsp14 activity was monitored.

### Mechanism Of Action

IC_50_ values were determined at a fixed concentration of RNA substrate (0.25 μM; 5x*K*_*m*_) and varying concentrations of SAM (up to 2.5 µM/10x *K*_*m*_), and varying concentrations of RNA (up to 0.5 µM; 10x *K*_*m*_) at fixed 1.25 µM (5x*K*_*m*_) of SAM. Linear increase in IC_50_ values as the concentration of substrate is increased, indicated a competitive pattern of inhibition as described by ^79^.

### Assessment of covalent binding by LC-MS

To form the protein–ligand (‘1911) complex, Nsp14 was incubated with 20 molar excess of compound for 2hr. at room temperature (20°C) before adding MS running buffer (0.1% FA). The resulting samples were separated on a HPLC column with 5-95% acetonitrile in water as eluent. The MS data were analyzed using an Agilent LC/MSD Time-of-Flight Mass Spectrometer equipped with an electrospray ionization source.

For compounds acryl42 and acryl42-10, an aliquot of pure Nsp14 enzyme was thawed on ice, centrifuged for 10 min at 14,000 rpm, diafiltrated, and concentrated to 40 μM in 50 mM HEPES pH 8.0, 20 mM NaCl, 5 mM MgCl_2_, 1 mM TCEP. Then, 1 μM Nsp14 was incubated alone or in the presence of 500 μM acryl42, or 50 μM acryl42-10 at room temperature in 100 μL aliquots. Time points were taken at 0, 2, 4, 8, and 18 h. For each time point, 1μL of the mix was injected into a Xevo G2-XS QTof Quadrupole Time of Flight mass spectrometer (Waters) using a solution of 0.05% formic acid at room temperature. Collected spectra from 700 to 1400 m/z were used to determine protein mass using MaxEnt with a 58500 to 62000 Da range and 1 Da/channel resolution.

### Compound Selectivity

Selectivity assays were performed as previously described ^80^. Compounds were tested at 10 μM in triplicate using radiometric assays. Enzymes with >50% inhibition were prioritized for concentration-response curves for IC_50_ determination.

### Aggregation

#### Dynamic Light Scattering (DLS)

Samples were prepared as 8-point half-log dilutions in filtered 50 mM KPi buffer, pH 7.0 with final DMSO concentration at 1% (v/v). Colloidal particle formation was detected using DynaPro Plate Reader II (Wyatt Technologies). All compounds were screened in triplicate at each concentration. For compounds that formed colloidal-like particles, the critical aggregation concentration (CAC) was determined by splitting the data into two data sets based on aggregating (i.e. >106 scattering intensity) and non-aggregating (i.e. <106 scattering intensity) and were fitted with separate nonlinear regression curves, and the point of intersection was determined using GraphPad Prism software version 9.1.1 (San Diego, CA).

#### Enzyme Inhibition Assays

Enzyme inhibition assays were performed at room temperature using CLARIOstar Plate Reader (BMG Labtech). Samples were prepared in 50 mM KPi buffer, pH 7.0 with final DMSO concentration at 1% (v/v). Compounds were incubated with 2 nM AmpC β-lactamase (AmpC) or Malate dehydrogenase (MDH) for 5 minutes. AmpC reactions were initiated by the addition of 50 μM CENTA chromogenic substrate or 50 μM Nitrocefin. The change in absorbance was monitored at 405 nm for CENTA(219475, Calbiochem) or 490 for Nitrocefin(484400, Sigma Aldrich) for 1 min 30 sec. MDH reactions were initiated by the addition of 200 μM nicotinamide adenine dinucleotide (NADH) (54839, Sigma Aldrich) and 200 μM oxaloacetic acid (324427, Sigma Aldrich). The change in absorbance was monitored at 340 nm for 1 min 30 sec. Initial rates were divided by the DMSO control rate to determine % enzyme activity. Each compound was screened at top concentration in triplicate. For compounds that showed greater than 40% inhibition against either enzyme, 8-point half-log concentration-response curves were performed in triplicate. Data was analyzed using GraphPad Prism software version 9.1.1 (San Diego, CA).

For detergent reversibility experiments, compounds that showed greater than 40% inhibition were screened again as 8-point half-log concentration-response curves in the presence of 0.01% (v/v) Triton X-100 in triplicates. Enzymatic reactions were performed/monitored as previously described.

#### UV-Vis Spectroscopy

Sample were prepared in 50mM KPi buffer, pH 7 with final DMSO concentration at 1% (v/v). Sample were loaded into 1.5mL methacrylate cuvette (14955128, FisherBrand) and measured using Cary UV-Vis Multicell Peltier (Agilent) from 200 nm to 800nm with a spectral bandwidth of 2nm. Data was analyzed using GraphPad Prism software version 9.1.1 (San Diego, CA).

### Statistical analyses

Data was analyzed using Prism 8.0 or 9.1.1 (GraphPad, San Diego, CA). For Nsp14 dose response curves, data was fitted to the four-parameter logistic equation.

## Supporting information

Singh_Nsp14_SI

## Data availability

The identities of compounds docked in this study are freely available from ZINC15, ZINC20, ZINC22, and covalent2022 databases (http://zinc15.docking.org, http://zinc20.docking.org, http://files.docking.org/zinc22, http://covalent2022.docking.org). Active compounds may be purchased from Enamine or WuXi or are available from the authors. All other data are available from the corresponding authors on request.

## Code availability

DOCK3.7 is freely available for non-commercial research (http://dock.compbio.ucsf.edu/DOCK3.7/), as is DOCKovalent. Web-based versions are freely usable by all (http://blaster.docking.org/). The ultra-large libraries used here are freely available (http://zinc15.docking.org, http://zinc20.docking.org, http://files.docking.org/zinc22).

## Acknowledgements

We thank OpenEye Software for Omega and additional tools, and Schrodinger LLC for the Maestro package. We thank Prof. Kevan Shokat and Dr. Qinheng Zheng for access to the Xevo G2-XS QTof Quadrupole Time of Flight mass spectrometer. We would like to also thank Structural Genomics Consortium (SGC) for their support. SGC is a registered charity (no: 1097737) that receives funds from Bayer AG, Boehringer Ingelheim, Bristol Myers Squibb, Genentech, Genome Canada through Ontario Genomics Institute (OGI-196), EU/EFPIA/OICR/McGill/KTH/Diamond Innovative Medicines Initiative 2 Joint Undertaking (EUbOPEN grant 875510), Janssen, Merck KGaA (EMD in Canada and US), Pfizer and Takeda.

## Funding

This work was supported by the US NIH R35GM122481 and DARPA HR0011-19-2-0020 (B.K.S.), NIH GM071896 and GM133836 (J.J.I.), and NIH U19AI171110 (D.G.F, B.K.S, M.V., J.J., M.O.). This work utilized the NMR Spectrometer Systems at Mount Sinai acquired with funding from National Institutes of Health SIG grants 1S10OD025132 and 1S10OD028504. M.O. received support from NIH U19AI171110 and the James B. Pendleton Charitable Trust.

## Author contributions

I.S. and E.A.F. performed docking screens, with input from B.K.S. Ligand optimization was conducted by I.S. and E.A.F. with input from B.K.S. E.A.F. and J.J.I. created the covalent libraries with assistance from X.W. F.L. performed Nsp14 enzymatic assays, Jump dilution and MOA experiments, and ‘**1911** LCMS. IC performed selectivity assays. A.L, and K.D. contributed to enzymatic assays. M.V., designed experiments, reviewed data and supervised. Additional LCMS testing was performed by A.R.-H. with supervision by D.G.F. M.R. and K.W. tested compounds in HeLa-ACE2 antiviral and cytotoxicity studies with supervision by A.G.-S. Cell viability in A549-ACE2 cells was performed by F.Z.B., supervised by M.O. Y.S.M. supervised Enamine compound synthesis purchased, assisted by N.A.T. J.J.I. built the ZINC15, ZINC22 ultra-large libraries. Synthesis of ZD160-68 was done by Z.D. with input and supervision by H.U.K. and J.J. B.K.S. and M.V. supervised the project with chemoinformatic input from J.J.I. I.S. and E.A.F. wrote the paper with input from the other authors and primary editing from M.V. and B.K.S. M.V. and B.K.S. conceived the project.

## Competing interests

B.K.S. serves on the SAB of Schrodinger and of Umbra Therapeutics, is a founder of Epiodyne, and with J.J.I of Deep Apple Therapeutics and BlueDolphin Leads LLC. The Jin laboratory received research funds from Celgene Corporation, Levo Therapeutics, Inc., Cullgen, Inc. and Cullinan Oncology, Inc. J.J. is a cofounder and equity shareholder in Cullgen, Inc. and a consultant for Cullgen, Inc., EpiCypher, Inc., and Accent Therapeutics, Inc.

## TOC Graphic

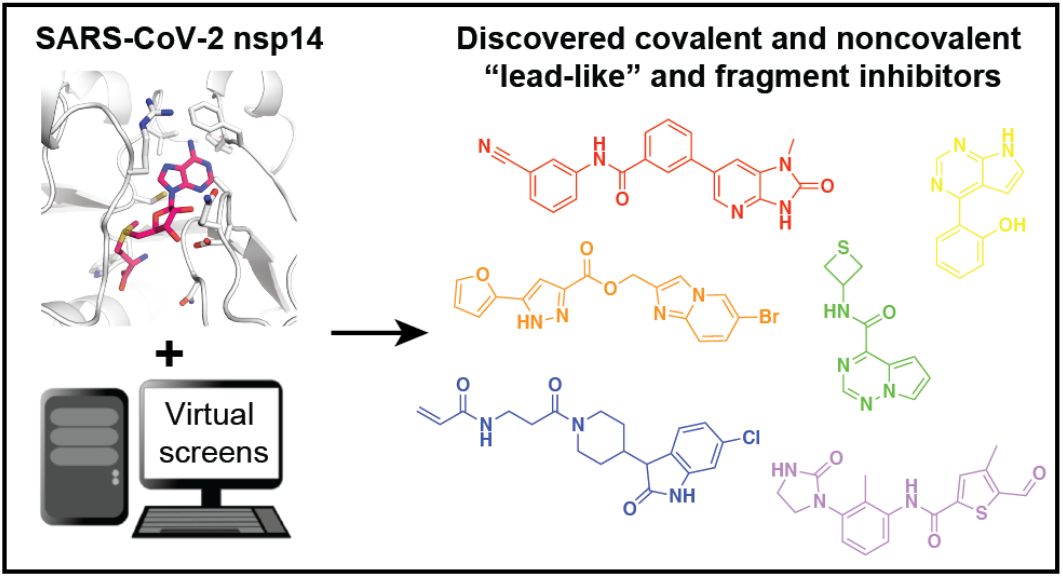

