## Supplementary material for "Structure-based discovery of inhibitors of the SARS-CoV-2 Nsp14 *N7*-methyltransferase": Singh_Nsp14_SI

**Contents:**

**Table S2.** Compound optimization for ‘2882.
**Table S3.** Compound optimization for ‘9213.
**Table S4.** Compound optimization for ‘4824.
**Table S5.** Summary of inhibitor aggregation testing.
**Table S6.** Compound optimization for acryl42.
**Table S7.** Docking inhibitors have novel, non-SAM like chemotypes.
**Supplementary Figure 1.** Assessment of reversibility of inhibition by rapid dilution.
**Supplementary Figure 2.** Mechanism of action of inhibitors.
**Supplementary Figure 3.1.** DLS scattering intensities for inhibitors.
**Supplementary Figure 3.2.** MDH inhibition in the absence and presence of Triton 0.01%.
**Supplementary Figure 3.3.** AmpC β-lactamase inhibition in the absence of Triton 0.01% with the nitrocefin substrate.
**Supplementary Figure 3.4.** Absorption spectrum for inhibitors.
**Supplementary Figure 4.** Concentration-response curves of non-covalent fragment hits.
**Supplementary Figure 5.** Docked poses of non-covalent fragment hits.
**Supplementary Figure 6.** Concentration-response curves of covalent hits.
**Supplementary Figure 7.** Docked poses of covalent hits.
**Supplementary Figure 8.** Evaluating possible covalent binding of compounds ‘1911, acryl42, and acryl42_10 to nsp14 by mass spectrometry.
**Supplementary Figure 9.** Analytical data for active compounds.

**Other Supplementary Material for this manuscript includes the following:**

**Table S1.** All compounds tested. (.xlsx)

**Table S2. Compound optimization for ‘2882.**

| 2D Chemical Structure | $IC_{50}$ (μM) | 2D Chemical Structure | $IC_{50}$ (μM) |
|---|---|---|---|
| 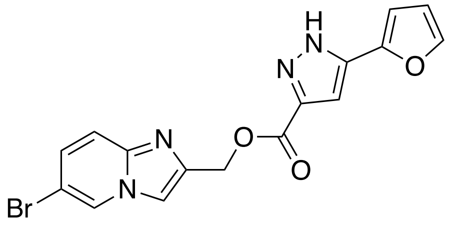 **Initial hit ZINC61142882 ('2882)** | **6** | 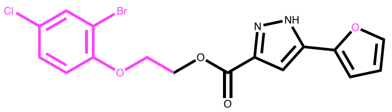 ZINC002325862871 (‘2817) | 18 |
| 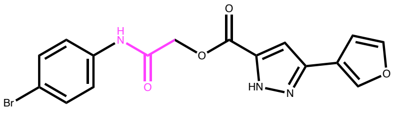 Z795161988 (‘1988) | 2.2 | 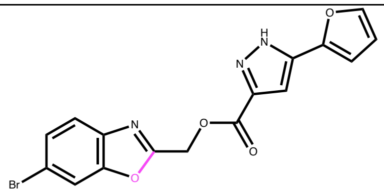 Z1724303092 (‘3092) | 21 |
| 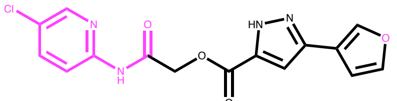 ZINC000919327864 (‘7864) | 7 | 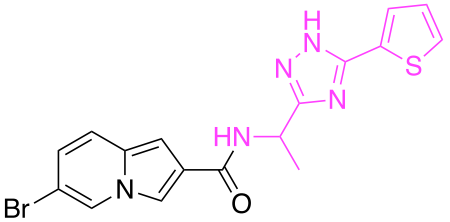 Z5347191013 (‘1013) | 30 |
| 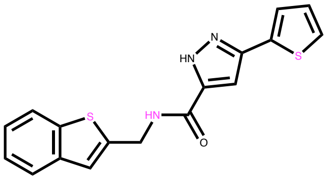 ZINC001141402437 (‘2437) | 15 | 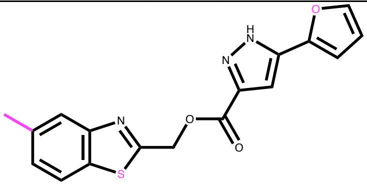 ZINC002325862682 (‘2682) | 38 |
| 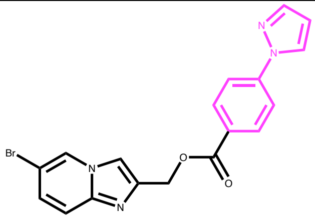 ZINC000918899102 (‘9102) | 17 | 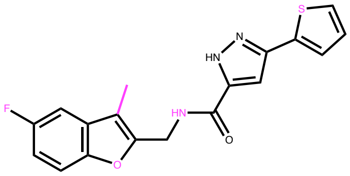 ZINC001444146032 (‘6032) | 28 |

**Table S3. Compound optimization for '9213.**

| 2D Chemical Structure | $IC_{50}$ (μM) | 2D Chemical Structure | $IC_{50}$ (μM) |
|---|---|---|---|
| 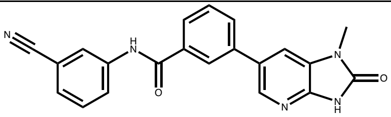<br>**Initial hit ZINC475239213 ('9213)** | **20** | 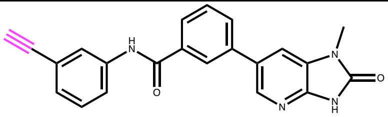<br>ZINC771823888 ('3888) | 27 |
| 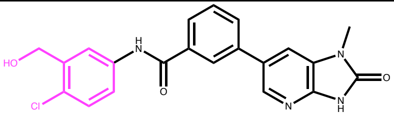<br>Z5347169163 ('9163) | 15 | 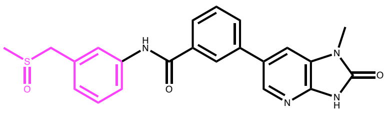<br>ZINC000475232808 ('2808) | 31 |
| 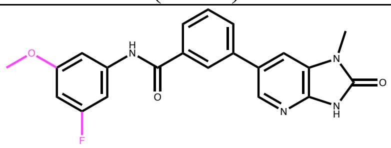<br>ZINC001342858621 ('8621) | 19 | 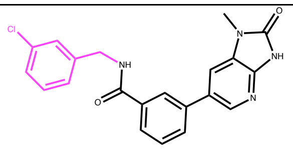<br>ZINC475241593 ('1593) | 44 |
| 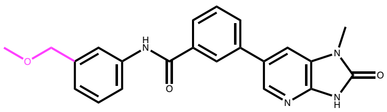<br>ZINC000475240670 ('0670) | 20 | 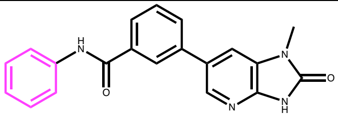<br>ZINC000355153196 ('3196) | 47 |

**Table S4. Compound optimization for ‘4824.**

| 2D Chemical Structure | $IC_{50}$ (μM) | 2D Chemical Structure | $IC_{50}$ (μM) |
|---|---|---|---|
| 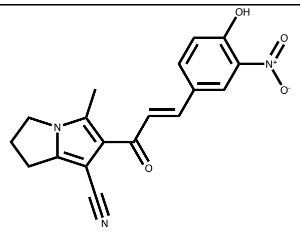<br>**Initial hit**<br>**ZINC730084824**<br>**('4824)** | **43** | 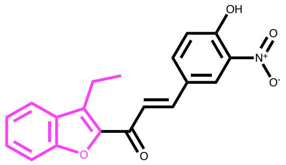<br>ZINC000916131631<br>(‘1631) | 25 |
| 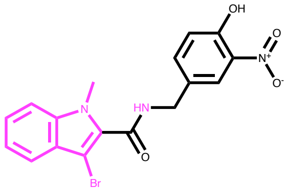<br>Z5347186947<br>(‘6947) | 19 | 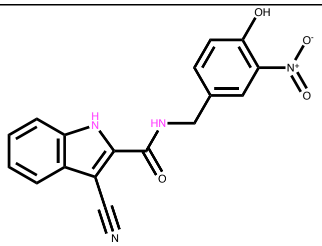<br>Z5347186943<br>(‘6943) | 40 |
| 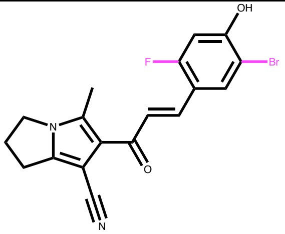<br>Z5347186953<br>(‘6953) | 22 | | |

### Table S5. Summary of inhibitor aggregation testing.

| Compound | DLS[a] CAC [µM] | MDH[b] $IC_{50}$ [µM] | AmpC[c] $IC_{50}$ [µM] | AmpC[d] $IC_{50}$ [µM] | Detergent reversibility[e] | Ab(340)[f] | Ab(405)[g] | Ab(490)[h] |
|---|---|---|---|---|---|---|---|---|
| **Acryl41** | no | no | no | NA | NA | yes | no | no |
| **Acryl42** | yes 20.2 | no | no | NA | NA | no | no | no |
| **‘4824** | no | yes 9.64 | yes NA | yes 35.1 | yes | yes | yes | slight |
| **‘6947 (‘4824 analog)** | yes 13.9 | yes 31.8 | no | NA | yes | yes | slight | slight |
| **‘6953 (‘4824 analog)** | yes 31.6 | yes 10.7 | yes NA | yes 35.1 | yes | yes | yes | no |
| **‘6943 (‘4824 analog)** | yes 10.5 | yes 37.2 | no | NA | yes | slight | slight | no |
| **‘1988 (‘2882 analog)** | yes 27.9 | yes 5.13 | yes NA | yes 54.7 | yes | no | no | no |
| **‘3888 (‘9213 analog)** | yes 10.5 | no | no | NA | NA | yes | no | no |
| **‘1911** | yes 4.41 | yes 17.8 | yes NA | no | yes | yes | slight | no |
| **‘1889** | yes 54.7 | no | no | NA | NA | no | no | no |
| **‘0222** | yes 112.2 | no | no | NA | NA | no | no | no |
| **‘0626** | no | no | no | NA | NA | slight | no | no |
| **‘0683** | no | yes 73.8 | yes NA | no | yes | no | no | no |
| **‘0741** | no | yes 91.6 | no | NA | no | no | no | no |
| **‘0772** | no | no | no | NA | NA | yes | no | no |
| **‘5604** | yes 37.2 | yes 94.4 | no | NA | yes | no | no | no |
| **‘5763** | yes 56.2 | no | no | NA | NA | no | no | no |
| **‘9744** | yes 168.7 | no | no | NA | NA | no | no | no |

[a] Scattering intensity measured with Dynamic Light Scattering (DLS). Yes = > $10^6$ scattering intensities; no = < $10^6$ scattering intensities. Critical aggregation concentrations (CACs) determined if applicable. Experiments were performed in triplicate.
[b] Enzymatic inhibition of MDH with cofactor NADH (absorption at 340 nm). Yes = inhibitor of MDH; no = does not inhibit MDH. $IC_{50}$ values included if applicable. Experiments were performed in triplicate.
[c] Enzymatic inhibition of AmpC β-lactamase with the substrate CENTA (absorption at 405 nm). Yes = inhibitor at 100 µM single point evaluation. Experiments were performed in triplicate.
[d] Enzymatic inhibition of AmpC β-lactamase with the substrate Nitrocefin (absorption at 490 nm). Yes = inhibitor of AmpC; no = does not inhibit AmpC. $IC_{50}$ values included if applicable. Experiments were performed in triplicate.
[e] Reversibility of MDH inhibition with 0.01% Triton. Yes = reversible inhibition; no = inhibition not reversed with addition of triton. Experiments were performed in triplicate.
[f] Compound absorption around λ=340 nm similar to MDH cofactor, NADH. Yes = high absorption, slight = some absorption, no = no absorption at this wavelength.
[g] Compound absorption around λ=405 nm similar to AmpC β-lactamase substrate, CENTA. Yes = high absorption, slight = some absorption, no = no absorption at this wavelength.
[h] Compound absorption around λ=490 nm similar to MDH substrate, Nitrocefin. Yes = high absorption, slight = some absorption, no = no absorption at this wavelength.
NA, not applicable
Red indicates a hit for the respective experiment. Yellow indicates a partial hit for the respective experiment.

**Table S6. Compound optimization for acryl42.**

| 2D Chemical Structure | $IC_{50}$ (μM) | 2D Chemical Structure | $IC_{50}$ (μM) |
|---|---|---|---|
| 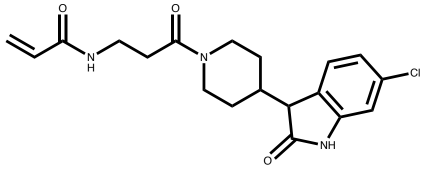 **Acryl42** | **32** | 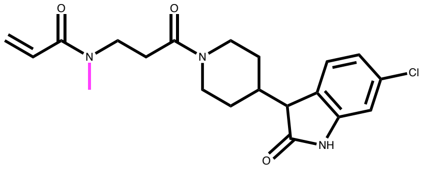 Acryl42_5 | NI |
| 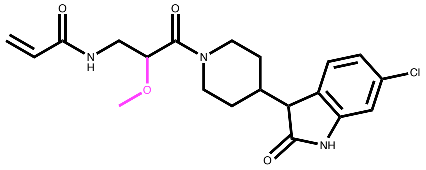 Acryl42_10 | 7 | 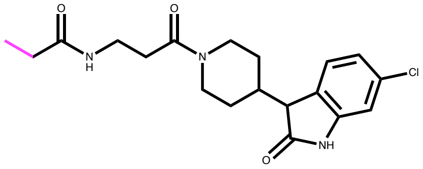 ZD160-68 | NI |
| 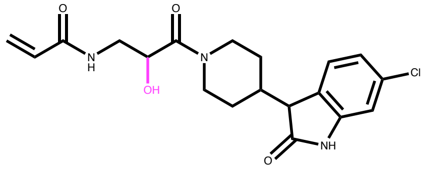 Acryl42_11 | >200 | | |

NI, no inhibition

**Table S7. Docking inhibitors have novel, non-SAM like chemotypes.**

| Compound name | $IC_{50}$ (μM) | 2D chemical structure | Molecular Weight<br>cLogP<br># charged groups | Similarity to SAM (TC) |
|---|---|---|---|---|
| **ZINC61142882 ('2882)** | **6** | 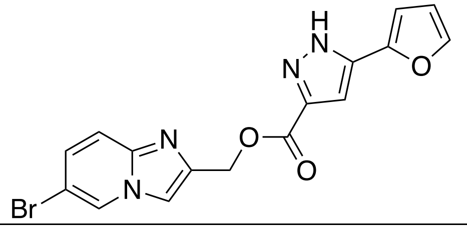 | 386.0<br>3.44<br>0 | 0.09 |
| Z795161988 | 2.2 | 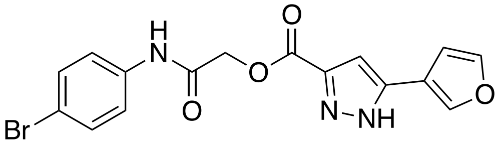 | 389.0<br>3.23<br>0 | 0.09 |
| **ZINC475239213 ('9213)** | **20** | 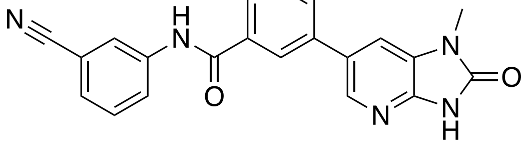 | 369.12<br>3.05<br>0 | 0.09 |
| Z5347169163 | 15 | 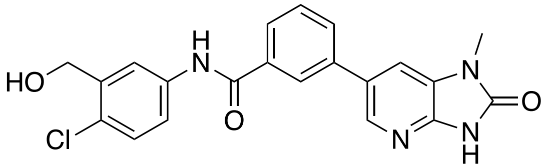 | 408.10<br>3.33<br>0 | 0.11 |
| ZINC001342858621 | 19 | 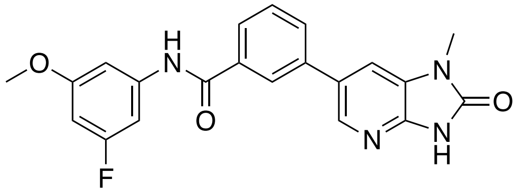 | 392.13<br>3.33<br>0 | 0.09 |
| **ZINC730084824 ('4824)** | **43** | 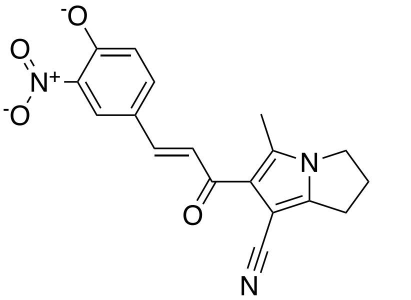 | 336.11<br>2.49<br>3 | 0.08 |
| ZINC000916131631 | 25 | 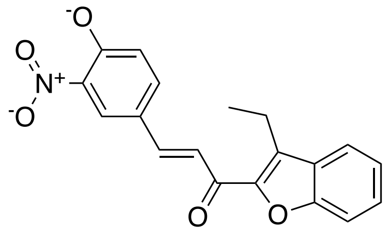 | 336.09<br>3.87<br>3 | 0.11 |

| | | | | |
|---|---|---|---|---|
| Z5347186953 | 22 | 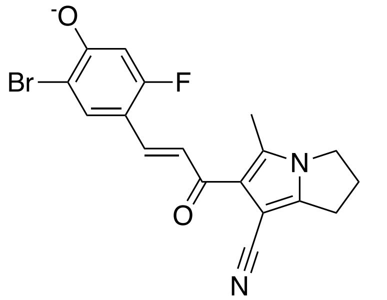 | 387.01<br>3.49<br>1 | 0.09 |
| Z5347186947 | 19 | 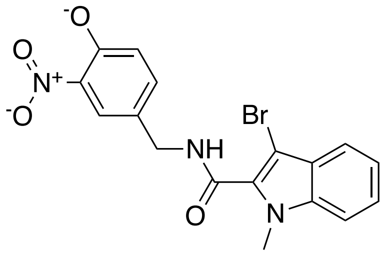 | 402.01<br>2.85<br>3 | 0.12 |
| Z5347186943 | 40 | 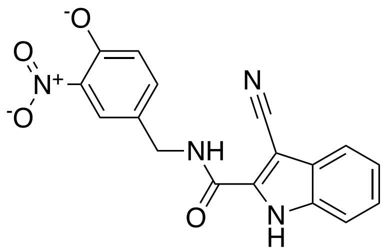 | 335.08<br>1.95<br>3 | 0.09 |
| ‘0683 | 12 | 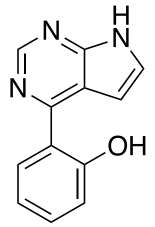 | 221.07<br>2.33<br>0 | 0.10 |
| ‘0626 | 60 | 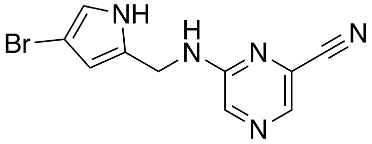 | 277.00<br>2.05<br>0 | 0.10 |
| ‘5763 | 62 | 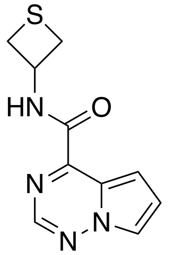 | 234.06<br>0.57<br>0 | 0.04 |
| ‘0772 | 78 | 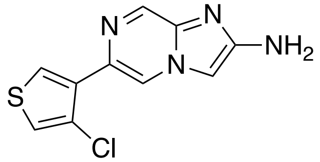 | 250.01<br>2.69<br>0 | 0.10 |

| | | | | |
|---|---|---|---|---|
| ‘4975 | 3.2 | 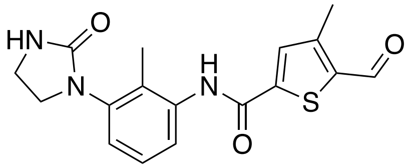 | 343.10<br>2.96<br>0 | 0.09 |
| ‘1911 | 3.8 | 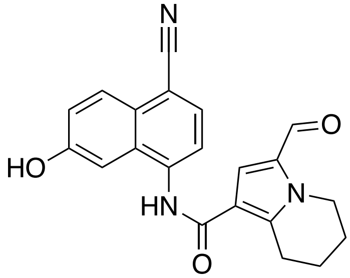 | 359.13<br>3.62<br>0 | 0.08 |
| ‘1889 | 12 | 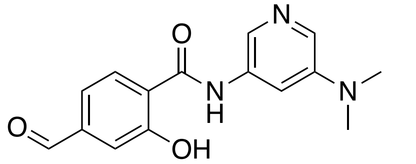 | 285.11<br>1.92<br>0 | 0.10 |
| acryl42 | 32 | 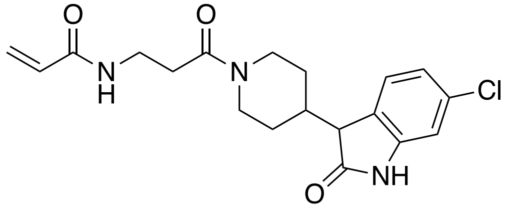 | 375.13<br>2.31<br>0 | 0.09 |
| acryl41 | 39 | 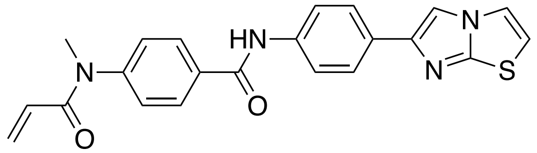 | 402.12<br>4.46<br>0 | 0.08 |
| ‘1903 | 19 | 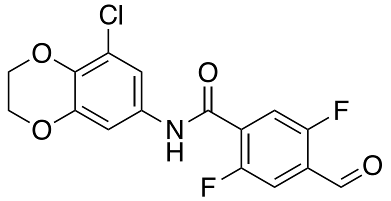 | 353.03<br>3.45<br>0 | 0.10 |
| acryl42-10 | 7 | 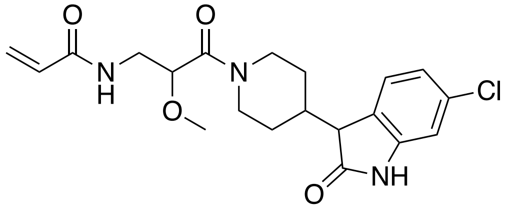 | 405.15<br>1.93<br>0 | 0.12 |

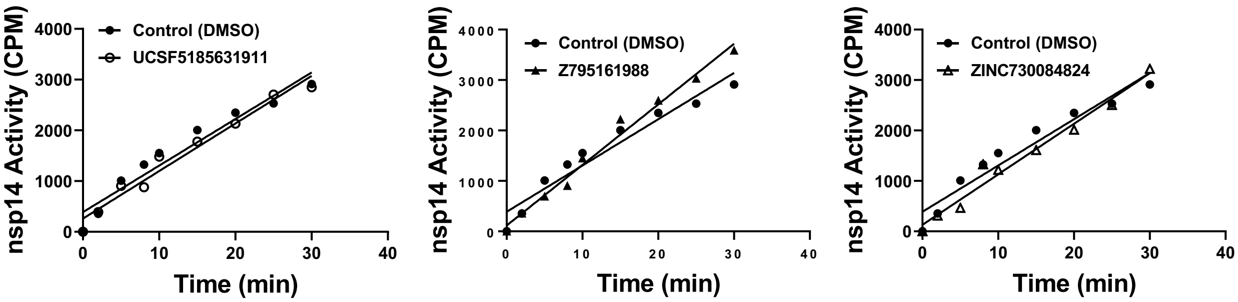


**Supplementary Figure 1. Assessment of reversibility of inhibition by rapid dilution.** The recovery of nsp14 activity after incubation with each inhibitor **(‘1911**, **‘1988**, **‘4824**) and rapid dilution was monitored. Nsp14 at 100-fold higher concentration than what required for activity measurement (1.5 nM) was incubated with each compound at 10-fold of $IC_{50}$ value concentration for 1h at room temperature. Reaction mixtures were then rapidly diluted 100-fold into the assay buffer containing substrate RNA and SAM, and recovery of the nsp14 activity was monitored.

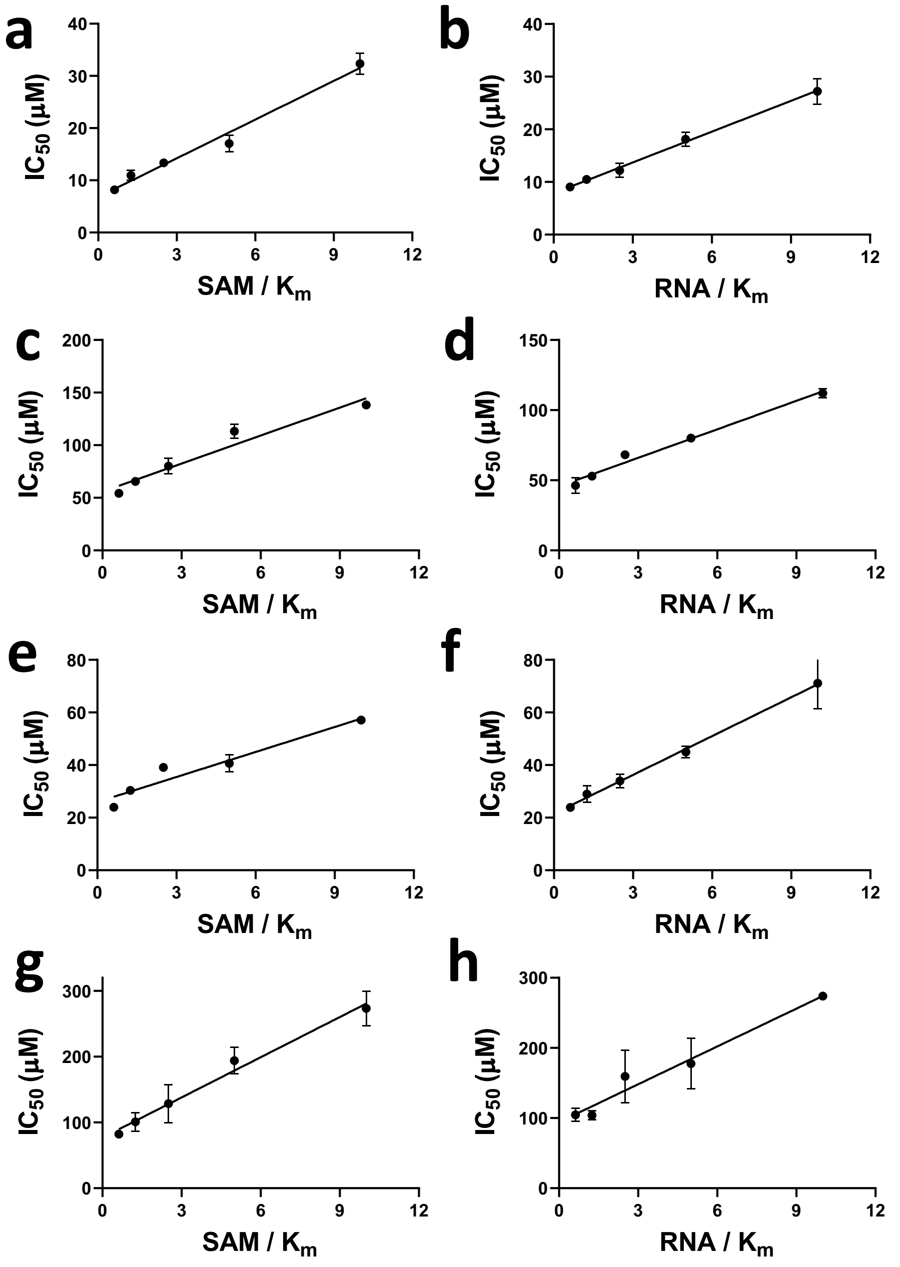


**Supplementary Figure 2. Mechanism of action of inhibitors. Compounds (a, b) '1988, (c, d) '4824, (e, f) '1911,** and **(g, h) '0683** were evaluated for their inhibition of SAM and RNA binding. **(a, c, e, g)** $IC_{50}$ values were determined at a fixed concentration of RNA substrate (0.25 µM; 5x$K_m$ ) and varying concentrations of SAM (up to 2.5 µM/10x $K_m$) and **(b, d, f, h)** varying concentrations of RNA (up to 0.5 µM; 10x $K_m$) and fixed 1.25 µM (5x$K_m$) of SAM. The experiments were performed in triplicate. Linear increase in $IC_{50}$ values as the concentration of substrate is increased, indicated a competitive pattern of inhibition.

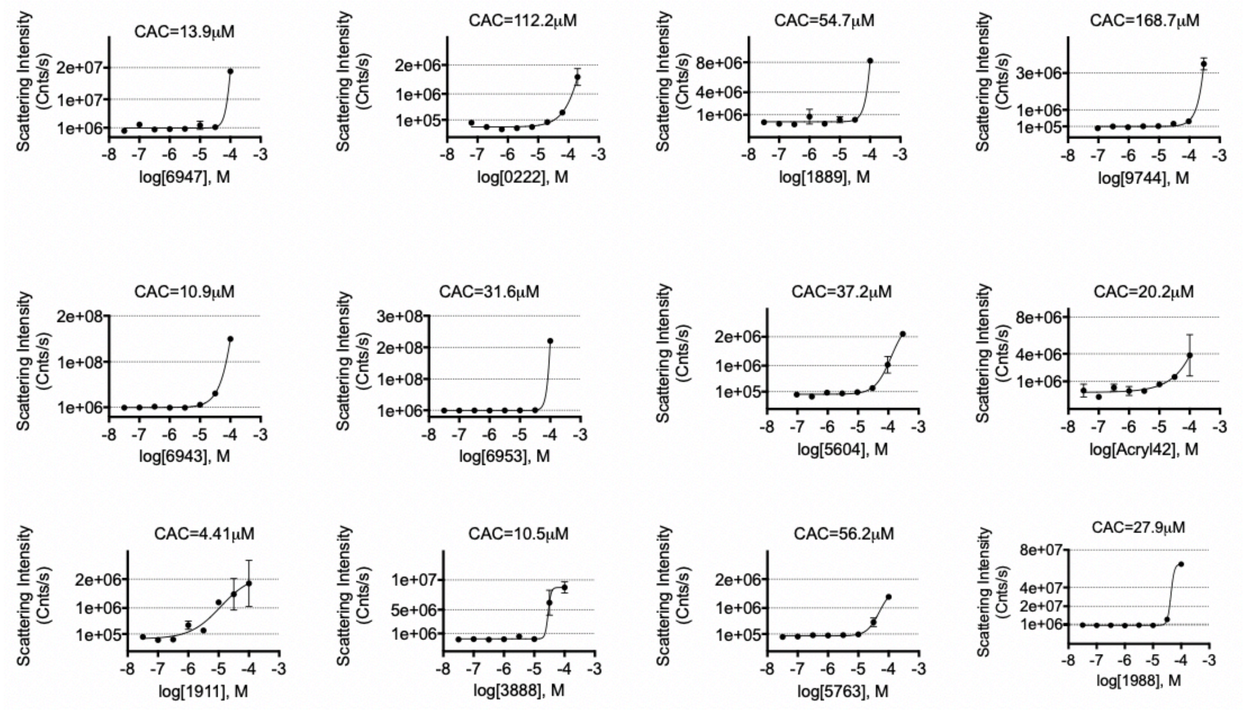


**Supplementary Figure 3.1.** DLS scattering intensities for inhibitors. Compounds with scattering intensity > 106 were further evaluated to establish CACs. Experiments were performed with n= 2 to 3. Results are summarized in Supplementary Table S5.

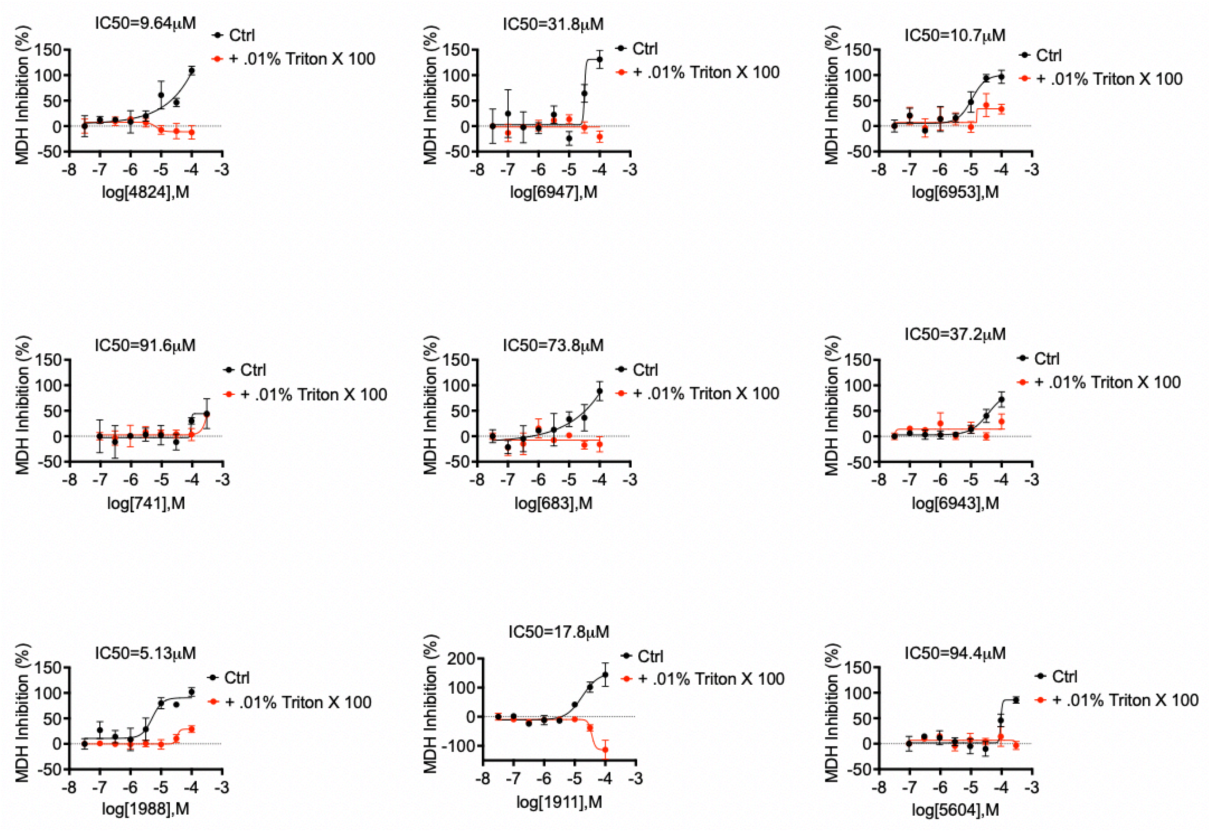


**Supplementary Figure 3.2. MDH inhibition in the absence and presence of Triton 0.01%.** Experiments were performed with n= 2 to 3. Results are summarized in Supplementary Table S5.

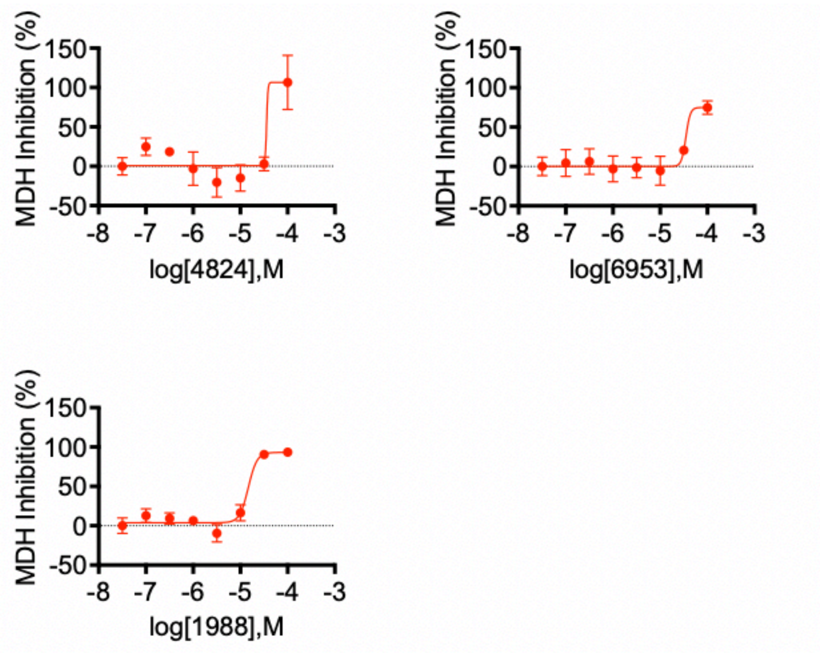


**Supplementary Figure 3.3. AmpC β-lactamase inhibition in the absence of Triton 0.01% with the nitrocefin substrate.** Experiments were performed with n= 2 to 3. Results are summarized in Supplementary Table S5.

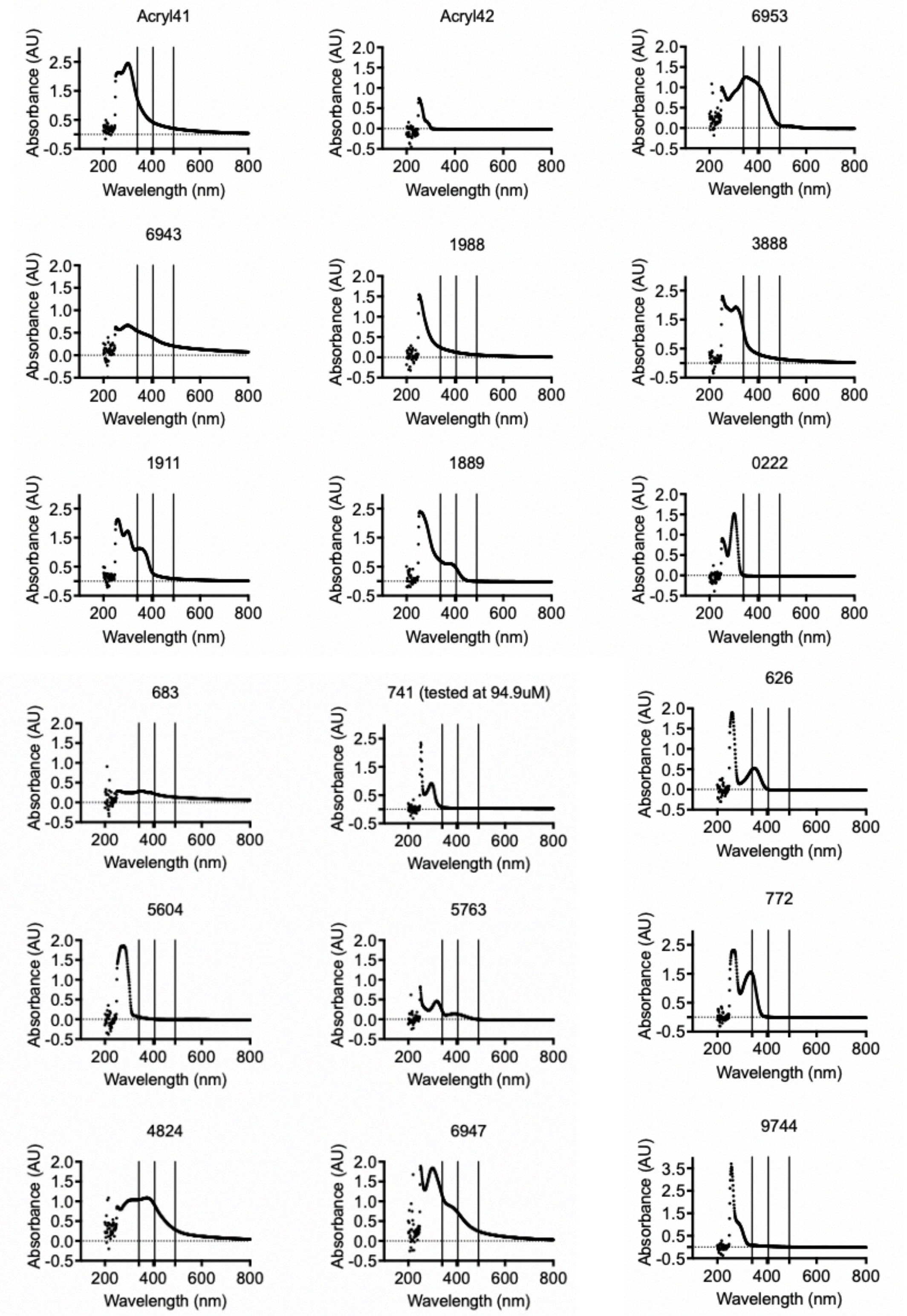
Acryl41
Acryl42
6953
6943
1988
3888
1911
1889
0222
683
741 (tested at 94.9uM)
626
5604
5763
772
4824
6947
9744
Absorbance (AU)
Wavelength (nm)

**Supplementary Figure 3.4. Absorption spectrum for inhibitors.** Compound absorption spectrum were determined to evaluate potential interference at enzymatic cofactor or substrate wavelengths. Vertical lines on the spectrum correspond to MDH cofactor NADH, AmpC β-lactamase CENTA, and AmpC β-lactamase Nitrocefin, respectively. Results are summarized in Supplementary Table S5.

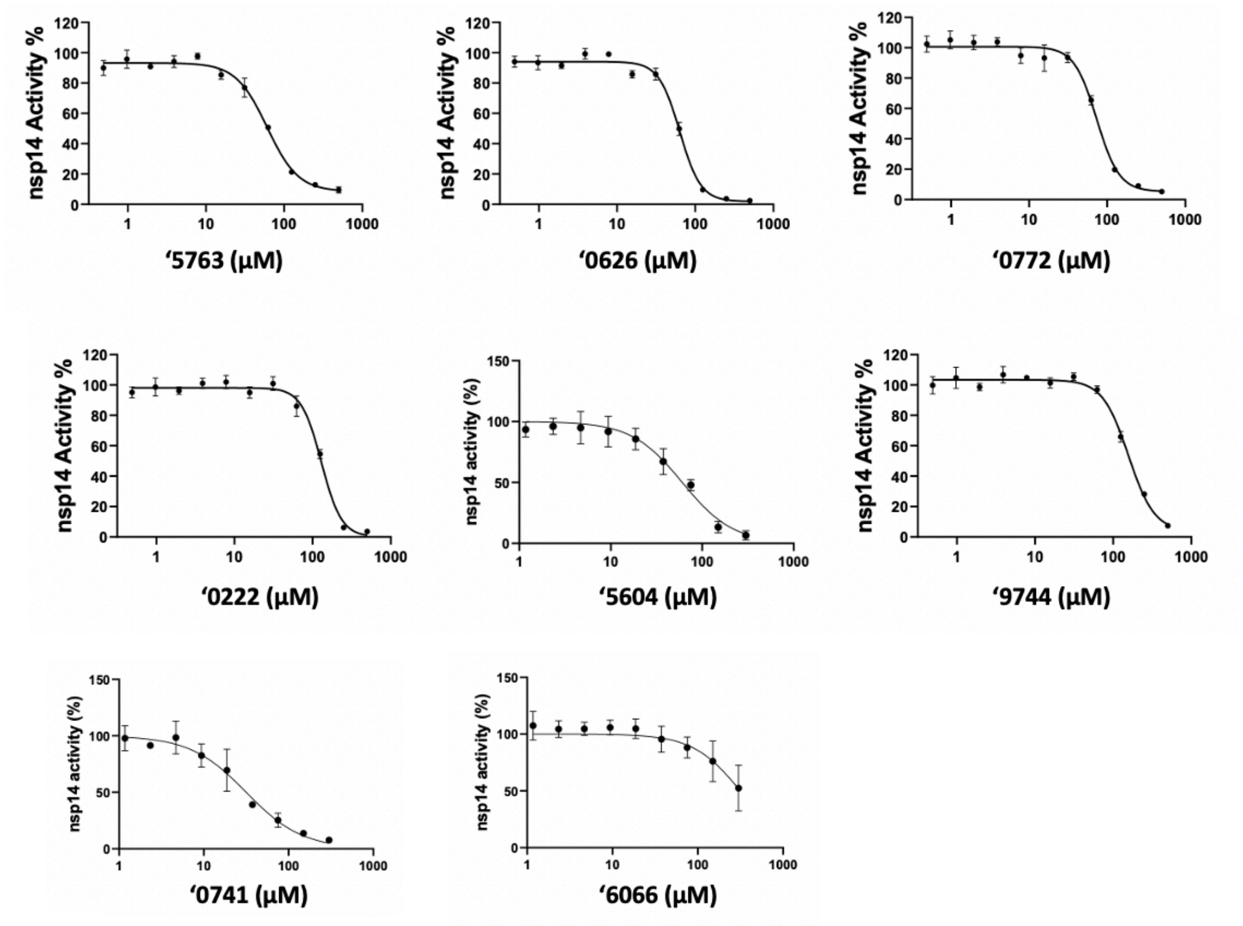


**Supplementary Figure 4. Concentration-response curves of non-covalent fragment hits.** Docking hits shown with their respective concentration-response curves in the *N7*-MTase inhibitory activity assay and docked poses. The experiments were performed in triplicate.

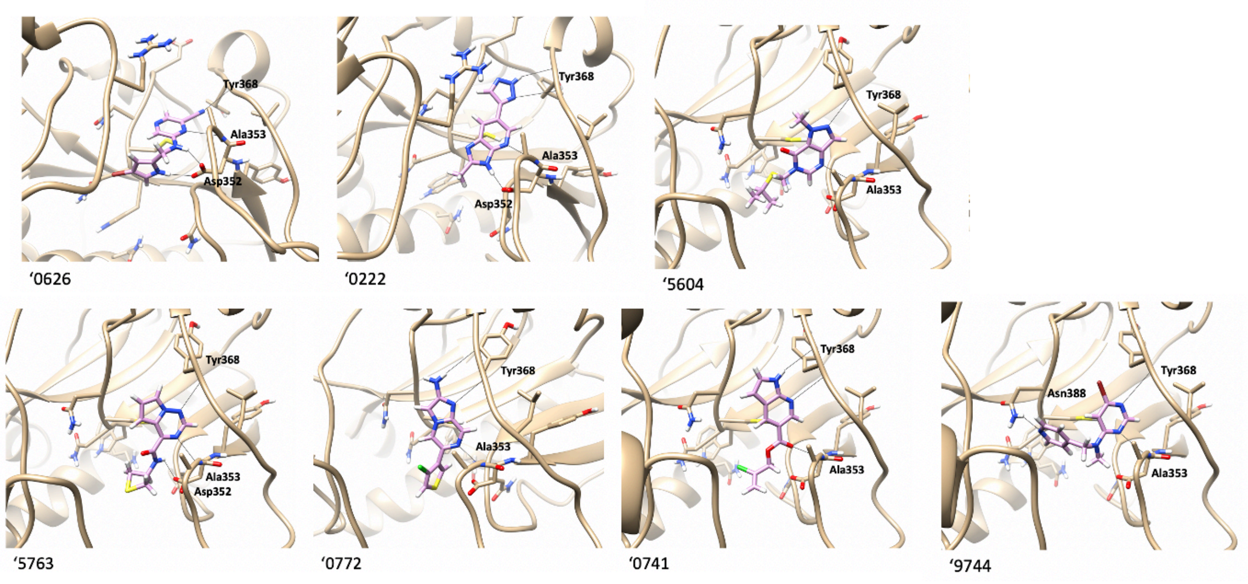


**Supplementary Figure 5. Docked poses of non-covalent fragment hits.**

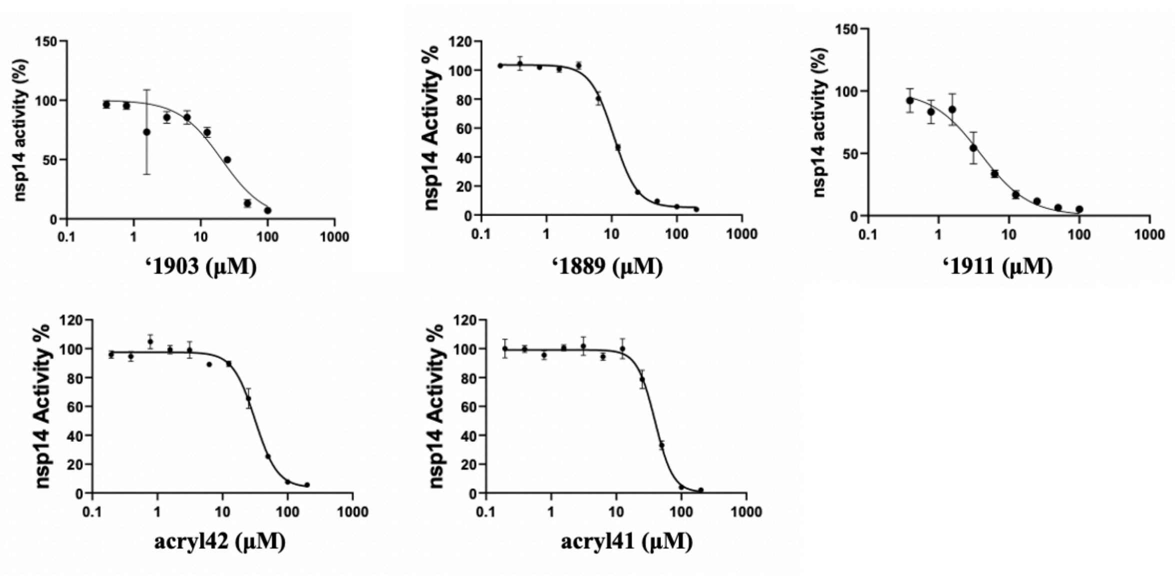


**Supplementary Figure 6. Concentration-response curves of covalent hits.** Docking hits shown with their respective concentration-response curves in the *N7*-MTase inhibitory activity assay and docked poses. The experiments were performed in triplicate.

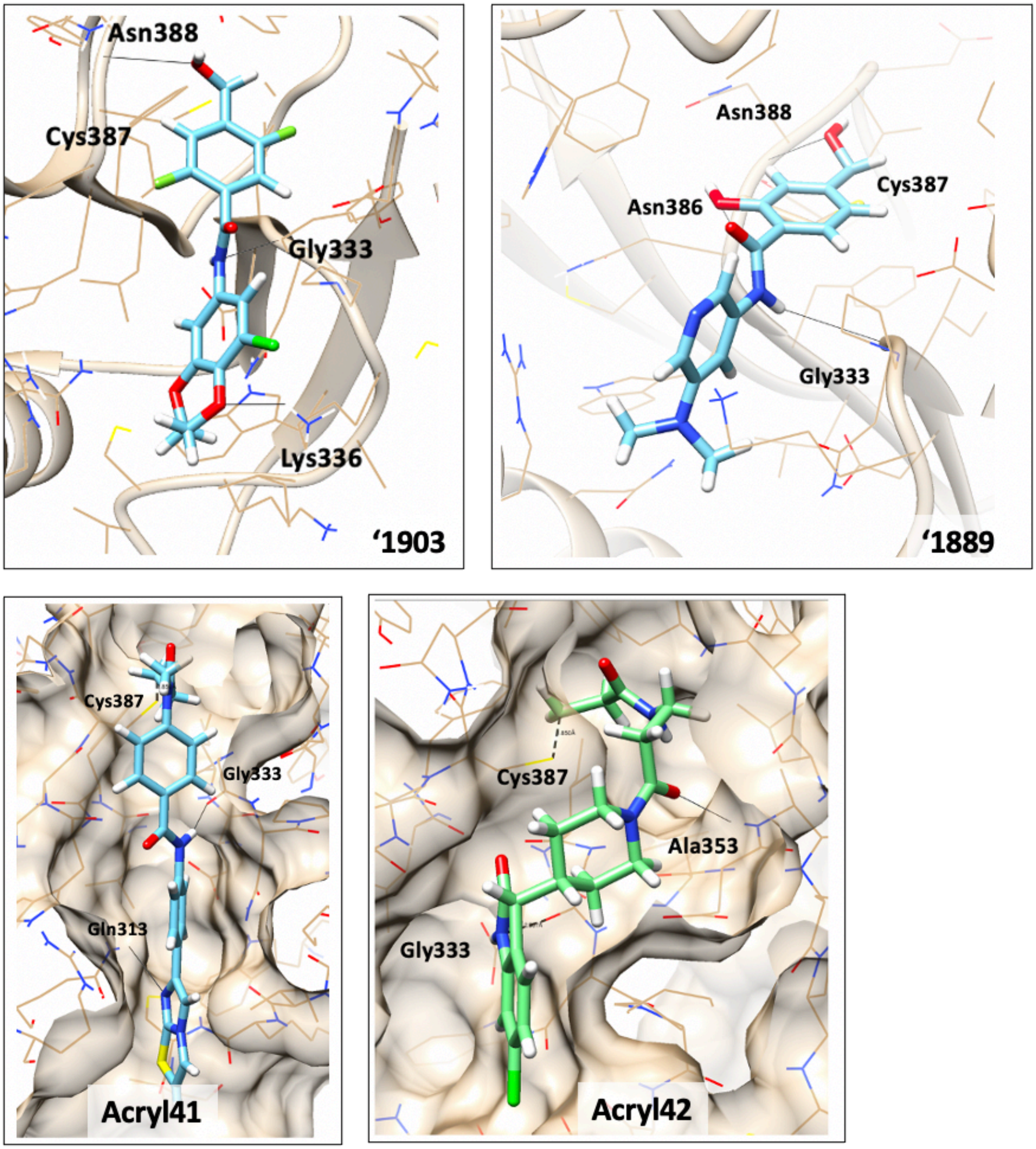


**Supplementary Figure 7. Docked poses of covalent hits.**

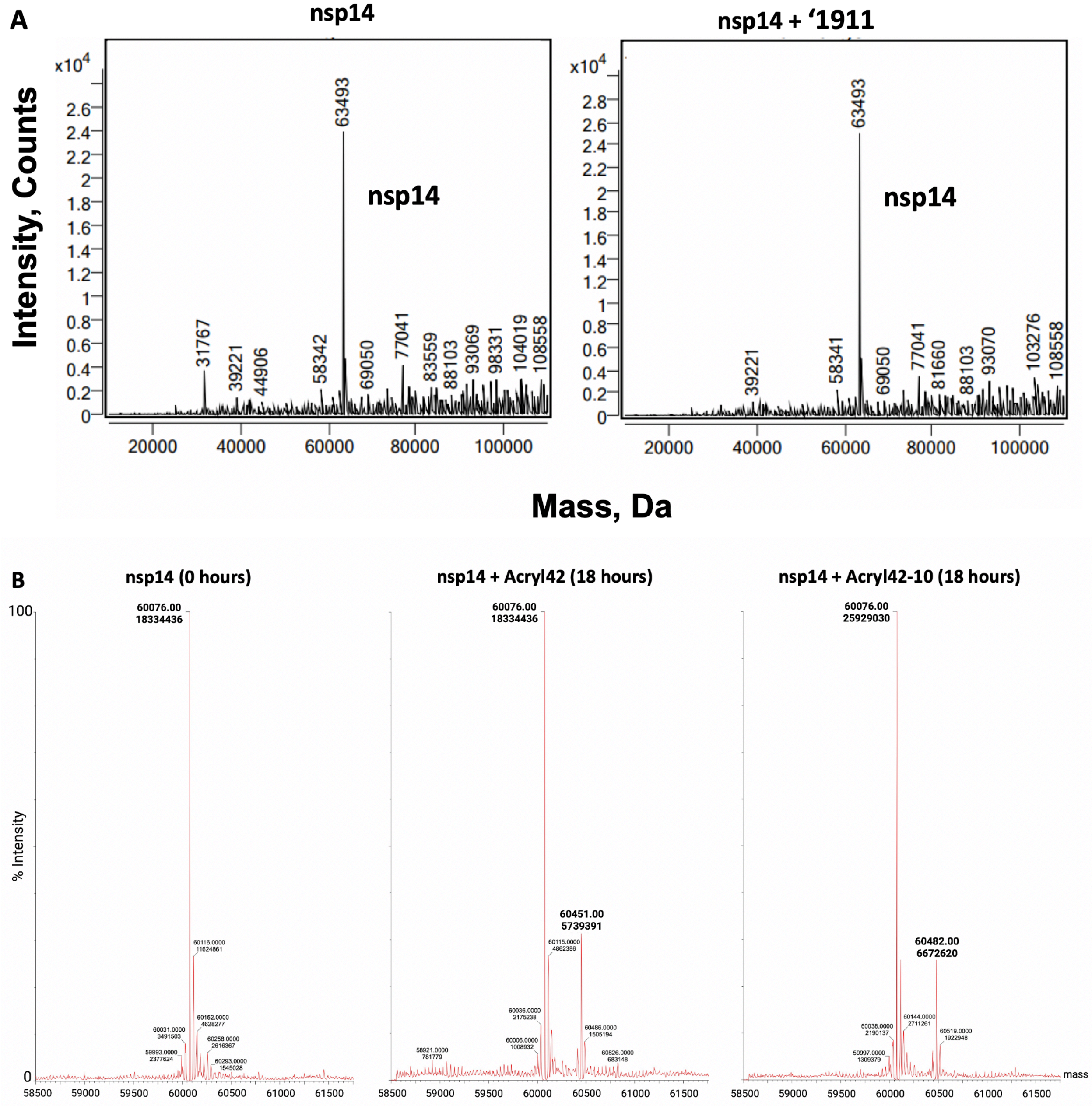


**Supplementary Figure 8. Evaluating possible covalent binding of compound ‘1911, acryl42, and acryl42-10 to nsp14 by mass spectrometry.** (**A**) Nsp14 was incubated in the absence (left) and presence (right) of compound 1911 at 20x molar excess for 2 hours at room temperature (22±1 ºC). Both samples were analyzed by mass spectrometry. No change in molecular mass of nsp14 upon incubation with compound 1911 confirmed that it is not an irreversible covalent inhibitor. (**B**) LC-MS observed mass and peak intensity data of nsp14 with no compound (left), in the presence of 500 µM acryl42 (middle), or 50 µM acryl42-10 (right). The peak at 60451 corresponds to nsp14 adduct with acryl42 (middle; fraction adduct 0.238) and the 60482 peak corresponds to the adduct of nsp14 with acryl42-10 (right; fraction adduct 0.205) both incubated at room temperature for 18 hours.

**Supplementary Figure 9. Analytical data for active compounds.**

**General synthetic procedures.**

**Method 1.** An amine (100 mg) and DIPEA (1.1 mol. eq. to the amine; additional equivalents were added when amine used was in salt form) were dissolved in 0.5 ml DMSO. The vial was then shaken for 20 min at room temperature (RT). Alkyl chloride (1 mol. eq. to the amine) was then added. The vial was sealed and stirred for 1 hour. Then the solution was heated for 8 hours at 100°C. After cooling down the mixture was filtered; the solvent and volatile components were evaporated under reduced pressure to give the crude product. The product was further purified by HPLC.

**Method 2**. Methylene-active compound (100 mg), aldehyde (1 mol. eq. to the methylene-active compound), DMF (0.5 ml), and trimethylchlorosilane (2.1 mol. eq. to the methylene-active compound) were placed in the vial. The vial was then placed in the thermostat (set to 100°C) for 48 hours. After cooling the reaction mixture down to RT, DIPEA (0.2 ml) was added to the vial, and it was stirred for 30 min. The solvents were evaporated under reduced pressure to give the crude product. The product was further purified by HPLC.

**Method 3.** An amine (100 mg), an acid (1.1 mol. eq. to the amine), and 0.5 ml of DMSO were placed into a 4 ml capped glass vial and the mixture was stirred for 30 min at RT. Then 1-Ethyl-3-(3-dimethylaminopropyl)carbodiimide (EDC, 1.2 mol. eq. to the amine) was added and the mixture was stirred for 1 hour. If the solution was transparent, the mixture was left overnight at room temperature as is; otherwise, the vial was placed in the ultrasonic bath and left overnight. The solution was filtered, and the solvent and volatile components were evaporated under reduced pressure to give the crude product. The product was further purified by HPLC.

**Method 4.** An amine (100 mg), DIPEA (1.2 mol. eq. to the amine), and DMSO (0.5 ml) were placed into a 4 ml capped glass vial and stirred for 30 min at RT. After the addition of an aryl halide (1.2 mol. eq. to the amine), the mixture was stirred for 1 hour at RT. Then the vial was placed on a thermostat (set to 100°C) for 9 hours. After cooling down the mixture was filtered; the solvent and volatile components were evaporated under reduced pressure to give the crude product. The product was further purified by HPLC.

**Method 5.** An aryl halide (100 mg), boronic derivative (1 mol. eq. to the aryl halide), sodium carbonate (1.5 mol. eq. to the aryl halide), 1,1′-[Bis(diphenylphosphino)ferrocene]-dichloropalladium(II) (catalytic amount) and water-dioxane mixture 1:3 (1 ml) were placed in a vial. The vial was then placed into a thermostat and stirred for 24 hours at 95°C. After cooling down the mixture, the solvents were evaporated under reduced pressure. The residue was dissolved in chloroform (3 ml) and washed with water (3x1 ml). The catalyst was then filtered, and solvents were evaporated under reduced pressure. The product was further purified by HPLC.

(6-bromoimidazo[1,2-a]pyridin-2-yl)methyl 5-(furan-2-yl)-1H-pyrazole-3-carboxylate – Z793205438 (Method 1).
Yield: 24%; purity, 94.2% (assesed by LC/MS).
$^{1}$H NMR (500 MHz, DMSO-$d_6$) δ 14.06 (s, 1H), 8.89 (s, 1H), 8.01 (s, 1H), 7.76 (s, 1H), 7.52 (d, $J$ = 9.5 Hz, 1H), 7.36 (dd, $J$ = 9.5, 1.8 Hz, 1H), 7.01 (s, 1H), 6.88 (s, 1H), 6.59 (s, 1H), 5.42 (s, 2H).
LC/MS (APSI) m/z [M+H] calculated for C16H12BrN4O3: 389.0; found: 389.0.

2-((4-bromophenyl)amino)-2-oxoethyl 3-(furan-3-yl)-1H-pyrazole-5-carboxylate – Z795161988 (Method 1).
Yield: 38%; purity, >95% (assesed by LC/MS).
$^1$H NMR (500 MHz, DMSO-$d_6$) δ 13.84 (s, 1H), 10.33 (s, 1H), 7.78 (s, 1H), 7.72 (s, 0H), 7.55 (d, $J$ = 8.6 Hz, 2H), 7.49 (d, $J$ = 8.7 Hz, 3H), 7.06 (s, 1H), 6.96 (s, 1H), 4.86 (s, 1H).
LC/MS (APSI) m/z [M+H] calculated for C16H13BrN3O4: 392.0; found: 392.0.

(E)-6-(3-(4-hydroxy-3-nitrophenyl)acryloyl)-5-methyl-2,3-dihydro-1H-pyrrolizine-7-carbonitrile – Z1143247121 (Method 2).
Yield: 29%; purity, >95% (assesed by LC/MS).
$^1$H NMR (400 MHz, DMSO-$d_6$) δ 11.54 (s, 1H), 8.22 (t, $J$ = 2.4 Hz, 1H), 7.90 (dt, $J$ = 8.6, 2.5 Hz, 1H), 7.56 (dd, $J$ = 15.7, 2.4 Hz, 1H), 7.42 (dd, $J$ = 15.6, 2.4 Hz, 1H), 7.19 (dd, $J$ = 8.7, 2.4 Hz, 1H), 4.06 – 3.96 (m, 2H), 3.01 – 2.92 (m, 2H), 2.46 (d, $J$ = 2.5 Hz, 3H).
LC/MS (APSI) m/z [M+H] calculated for C18H16N3O4: 338.1; found: 338.2.

N-(3-(4-(6-chloro-2-oxoindolin-3-yl)piperidin-1-yl)-3-oxopropyl)acrylamide – Z1479200718 (Method 3).
Yield: 35%; purity, 95% (assesed by LC/MS).
$^1$H NMR (500 MHz, DMSO-$d_6$) δ 10.50 (s, 1H), 8.11 – 8.01 (m, 1H), 7.22 (dd, $J$ = 8.0, 1.0 Hz, 1H), 6.96 (dt, $J$ = 8.0, 1.7 Hz, 1H), 6.80 (d, $J$ = 2.0 Hz, 1H), 6.02 (ddd, $J$ = 16.8, 13.7, 2.3 Hz, 1H), 4.43 (dd, $J$ = 21.4, 13.5 Hz, 1H), 3.81 (t, $J$ = 14.6 Hz, 1H), 3.46 – 3.41 (m, 1H), 3.27 (q, $J$ = 6.6 Hz, 2H), 2.90 (s, 1H), 2.43 (dq, $J$ = 12.8, 9.3, 8.7 Hz, 3H), 2.21 (td, $J$ = 12.3, 3.6 Hz, 1H), 1.56 (d, $J$ = 13.1 Hz, 1H), 1.43 (d, $J$ = 13.1 Hz, 1H).
LC/MS (APSI) m/z [M+H] calculated for C19H23ClN3O3: 376.1; found: 376.0.

N-(3-cyanophenyl)-3-(1-methyl-2-oxo-2,3-dihydro-1H-imidazo[4,5-b]pyridin-6-yl)benzamide – Z1723430981 (Method 3).
Yield: 21%; purity, >95% (assesed by LC/MS).
$^1$H NMR (500 MHz, DMSO-$d_6$) δ 11.59 (s, 1H), 10.63 (s, 1H), 8.32 (d, $J$ = 2.0 Hz, 1H), 8.25 (dt, $J$ = 3.7, 1.8 Hz, 2H), 8.06 (dt, $J$ = 7.3, 2.2 Hz, 1H), 7.97 – 7.89 (m, 2H), 7.83 (d, $J$ = 2.0 Hz, 1H), 7.64 (t, $J$ = 7.7 Hz, 1H), 7.62 – 7.54 (m, 2H), 3.36 (s, 3H).
LC/MS (APSI) m/z [M+H] calculated for C21H16N5O2: 370.1; found: 370.2.

5-(1H-indol-2-yl)-1-methyl-1H-pyrazol-4-amine – Z2732986066 (Method 5).
Yield: 28%; purity, >95% (assesed by LC/MS).
LC/MS (APSI) m/z [M+H] calculated for C12H13N4: 213.1; found: 213.2.

6-((isopropylthio)methyl)-1-methyl-1H-pyrazolo[4,3-d]pyrimidin-7(6H)-one – Z3343635604 (Method 1).
Yield: 29%; purity, >95% (assesed by LC/MS).
$^1$H NMR (500 MHz, DMSO-$d_6$) δ 8.26 (s, 1H), 7.98 (s, 1H), 5.16 (s, 2H), 4.19 (s, 3H), 3.09 (p, $J$ = 6.7 Hz, 1H), 1.21 (d, $J$ = 6.7 Hz, 6H).
LC/MS (APSI) m/z [M+H] calculated for C10H15N4OS: 239.1; found: 239.0.

N-(4-(imidazo[2,1-b]thiazol-6-yl)phenyl)-4-(N-methylacrylamido)benzamide – Z3756698609 (Method 3).

Yield: 51%; purity, >95% (assesed by LC/MS).
$^1$H NMR (500 MHz, DMSO-$d_6$) δ 10.33 (s, 1H), 8.16 (d, *J* = 2.7 Hz, 1H), 8.01 (dd, *J* = 8.7, 2.4 Hz, 2H), 7.92 (dd, *J* = 4.6, 2.6 Hz, 1H), 7.80 (s, 3H), 7.79 (d, *J* = 6.1 Hz, 1H), 7.47 – 7.40 (m, 2H), 7.23 (dd, *J* = 4.6, 2.5 Hz, 1H), 6.22 – 6.07 (m, 2H), 5.61 (dd, *J* = 9.6, 3.0 Hz, 1H), 3.30 (d, *J* = 4.3 Hz, 4H).
LC/MS (APSI) m/z [M+H] calculated for C22H19N4O2S: 403.1; found: 403.0.

N-(thietan-3-yl)pyrrolo[2,1-f][1,2,4]triazine-4-carboxamide – Z4324535763 (Method 3).
Yield: 26%; purity, >95% (assesed by LC/MS).
$^1$H NMR (500 MHz, DMSO-$d_6$) δ 9.70 (d, *J* = 8.8 Hz, 1H), 8.65 (d, *J* = 2.9 Hz, 1H), 8.24 (s, 1H), 7.44 (t, *J* = 3.8 Hz, 1H), 7.17 (q, *J* = 3.3 Hz, 1H), 3.71 – 3.62 (m, 2H), 3.30 (s, 1H), 3.19 (t, *J* = 8.1 Hz, 2H).
LC/MS (APSI) m/z [M+H] calculated for C10H11N4OS: 235.1; found: 235.0.

N-(5-(dimethylamino)pyridin-3-yl)-4-formyl-2-hydroxybenzamide – Z5185631889 (Method 3).
Yield: 58%; purity, 95% (assesed by LC/MS).
$^1$H NMR (500 MHz, DMSO-$d_6$) δ 11.44 (s, 1H), 9.95 (s, 1H), 8.18 (s, 1H), 7.96 (d, *J* = 7.9 Hz, 1H), 7.86 (d, *J* = 2.8 Hz, 1H), 7.54 (t, *J* = 2.4 Hz, 1H), 7.34 (d, *J* = 1.6 Hz, 1H), 7.33 – 7.27 (m, 1H), 2.92 (s, 7H).
LC/MS (APSI) m/z [M+H] calculated for C15H16N3O3: 286.1; found: 286.2.

2-methyl-6-(1H-1,2,3-triazol-4-yl)-1H-imidazo[4,5-b]pyridine – Z5348530222 (Method 5).
Yield: 34%; purity, >95% (assesed by LC/MS).
$^1$H NMR (500 MHz, DMSO-$d_6$) δ 8.83 (s, 1H), 8.33 (s, 1H), 2.58 (s, 3H).
LC/MS (APSI) m/z [M+H] calculated for C9H9N6: 201.1; found: 201.2.

6-(((4-bromo-1H-pyrrol-2-yl)methyl)amino)pyrazine-2-carbonitrile – Z5348530626 (Method 4).
Yield: 48%; purity, >95% (assesed by LC/MS).
$^1$H NMR (500 MHz, DMSO-$d_6$) δ 11.08 (s, 1H), 8.19 (d, *J* = 6.6 Hz, 2H), 8.02 (t, *J* = 5.3 Hz, 1H), 6.79 (dd, *J* = 2.8, 1.8 Hz, 1H), 6.06 – 6.00 (m, 1H), 4.33 (d, *J* = 5.2 Hz, 2H).
LC/MS (APSI) m/z [M+H] calculated for C10H9BrN5: 278.0; found: 278.0.

2-(7H-pyrrolo[2,3-d]pyrimidin-4-yl)phenol – Z5348530683 (Method 5).
Yield: 33%; purity, >95% (assesed by LC/MS).
$^1$H NMR (600 MHz, DMSO-$d_6$) δ 14.43 (s, 1H), 12.56 (s, 1H), 8.83 (s, 1H), 7.74 (d, *J* = 3.6 Hz, 1H), 7.43 – 7.37 (m, 1H), 7.12 (d, *J* = 3.6 Hz, 1H), 7.04 – 6.96 (m, 2H).
LC/MS (APSI) m/z [M+H] calculated for C12H10N3O: 212.1; found: 212.2.

2-chloroallyl 1H-pyrrolo[2,3-b]pyridine-5-carboxylate – Z5348530741 (Method 1).
Yield: 50%; purity, >95% (assesed by LC/MS).
$^1$H NMR (500 MHz, DMSO-$d_6$) δ 12.12 (s, 1H), 8.81 (d, *J* = 2.1 Hz, 1H), 8.57 (d, *J* = 2.3 Hz, 1H), 7.61 (dd, *J* = 3.5, 2.5 Hz, 1H), 6.63 (dd, *J* = 3.5, 1.9 Hz, 1H), 5.55 (d, *J* = 2.1 Hz, 1H), 4.96 (s, 2H).
LC/MS (APSI) m/z [M+H] calculated for C11H10ClN2O2: 237.0; found: 237.0.

6-(4-chlorothiophen-3-yl)imidazo[1,2-a]pyrazin-2-amine – Z5348530772 (Method 5).

Yield: 23%; purity, >95% (assesed by LC/MS).
$^{1}$H NMR (500 MHz, DMSO-$d_6$) δ 8.72 (t, *J* = 1.2 Hz, 1H), 8.57 (s, 1H), 7.94 (dd, *J* = 3.7, 1.1 Hz, 1H), 7.72 (dd, *J* = 3.6, 1.1 Hz, 1H), 7.26 (s, 1H), 5.62 (s, 2H).
LC/MS (APSI) m/z [M+H] calculated for C10H8ClN4S: 251.0; found: 251.0.
